# Intravital microscopy of satellite cell dynamics and their interaction with myeloid cells during skeletal muscle regeneration

**DOI:** 10.1101/2023.02.01.526724

**Authors:** Yingzhu He, Youshan Heng, Zhongya Qin, Xiuqing Wei, Zhenguo Wu, Jianan Qu

**Author notes:** These authors contributed equally to this work (Yingzhu He, Youshan Heng). Corresponding authors (J.Y.Q.); (Z.G.W).

## Abstract

Skeletal muscle regeneration requires the highly coordinated cooperation of muscle satellite cells (MuSCs) with other cellular components. However, due to technical limitations, it remains unclear how MuSCs dynamically interact with non-myogenic cells, especially myeloid cells, in live animals. In this work, we developed a dual-laser multimodal nonlinear optical microscope platform to serve as an effective tool for studying the real-time interaction between MuSCs and non-myogenic cells during the early phase of muscle regeneration. Increased cell volume and mitochondrial mass, cell density, and myotube formation are indicative of MuSCs activation/growth, proliferation, and differentiation, respectively. Using 3D time-lapse imaging on live reporter mice containing both red fluorescence protein (RFP)-labeled macrophages and yellow fluorescence protein (YFP)-labeled MuSCs, and taking advantages of the autofluorescence of reduced nicotinamide adenine dinucleotide (NADH), we monitored the real-time spatiotemporal interaction between RFP^+^ macrophages/RFP^-^ non-myogenic cells and YFP^+^ muscle stem/progenitor cells during the activation and the proliferation stages of regeneration. Our results indicated that their cell-cell contact was transient in nature. By inhibiting macrophage infiltration, we further showed that direct cell-cell contact between macrophages and MuSCs was not required for early activation of MuSCs before the proliferation stage. However, decreased macrophage infiltration impeded the proliferation and differentiation of MuSCs and also led to intramuscular fibrosis. Besides, neutrophil depletion in the CCR2 deficient mice did not delay the initial growth of MuSCs. These findings provide a new perspective on myeloid cells’ role during muscle regeneration.

## Introduction

Skeletal muscles, a type of striated muscle, are attached to bones and responsible for voluntary body movement. They have remarkable capacity for regeneration in response to injury^1^. The Paired Box 7 (Pax7)–expressing muscle stem cells (MuSCs; also known as satellite cells), residing between the sarcolemma and basal membrane of their host myofibers in quiescence^2,3^, are crucial for muscle homeostasis and regeneration. Upon injury or disease, satellite cells are activated with rapid expression of myogenic differentiation 1 (MyoD) protein and a gradual increase in cell size (also termed cell growth). The majority of activated MuSCs start to proliferate, differentiate into myocytes and finally fuse into existing injured fibers or fuse with each other to form new muscle fibers^3,4,5^. Moreover, muscle regeneration was abolished when adult Pax7^+^ satellite cells were inducibly depleted using a diphtheria toxin receptor (DTR) knock-in mouse model^6,7^.

Apart from satellite cells, non-myogenic cells, such as endothelial cells, fibroblasts, and immune cells, also participate in muscle regeneration by secreting cytokines to regulate the dynamics of satellite cells^3,7,8^. Among the immune cells, the myeloid lineage dominates the inflammatory infiltration at the early stage after muscle injury^8,9,10,11^. Macrophages are the predominant myeloid lineage cells observed during skeletal muscle regeneration, coordinating different biological processes^12^. Macrophage subtypes are primarily defined from *in vitro* studies by diverse environmental stimulation. Two main subtypes correspond to varying activation states and exhibit various functions. M1 macrophages (classically activated) exert pro-inflammatory activities and contribute to tissue damage; M2 macrophages (alternatively activated) are responsible for the resolution of inflammation and promote tissue regeneration^13^. In culture, M1 macrophages are responsible for the phagocytosis of muscle debris and promote the growth and proliferation of myoblasts, whereas M2 macrophages enhance the differentiation of myoblasts and the formation of myofibers^14^. *In vivo* analysis of macrophages using transgenic mice indicates that the CCR2^pos^CX3CR1^low^Ly6C^pos^ (pro-inflammatory) monocytes/macrophages (MOs/MΦs) infiltrate the injured sites mainly via the CCL2-CCR2 axis after acute skeletal muscle injury^12,15^. These MOs/MΦs convert to CCR2^neg/low^CX3CR1^hi^Ly6C^neg^ MOs/ MΦs *in situ* and then participate in the resolution of inflammation^14,15,16,17^. Based on the measurement of selected gene expression, CCR2^pos^CX3CR1^low^Ly6C^pos^ and CCR2^neg/low^CX3CR1^hi^Ly6C^neg^ macrophage subpopulations *in vivo* were considered equivalent to M1 and M2 macrophages, respectively. Nevertheless, the *in vitro*-defined M1/M2 paradigm was shown to be an inappropriate representation of macrophage heterogeneity during tissue regeneration *in vivo* because of the complex environmental stimulation^15,18^. Disrupting the CCL2-CCR2 axis by using CCR2- or CCL2- deficient mouse strains significantly reduced the recruitment of MOs/MΦs and impeded muscle regeneration^19,20,21^. Similarly, partial or total depletion of circulating monocytes leads to impaired muscle regeneration, as indicated by the smaller cross-sectional area of regenerating myofibers, intramuscular fat accumulation, fibrosis, or delayed angiogenesis in injured sites^14,19,22,23,24,25^.

Neutrophils, another prominent population of myeloid cells, are the earliest immune responders to infiltrate the damaged muscle within 1-3 hours after injury and are responsible for the establishment of necrotic microenvironment by releasing pro-inflammatory cytokines to attract other types of immune cells^11^. In one study, the depletion of neutrophils delayed muscle regeneration after injection of myotoxic substances in mice^26^. However, another study in mice showed that neutrophil depletion attenuated muscle injury after exhaustive exercise^27^. Inconsistency between these findings may be caused by different injury models. Besides, *in vitro* studies demonstrated that neutrophil-mediated factors promoted proliferation but impeded the differentiation of the myogenic progenitor cells reflecting a complex role for neutrophils in regulating myogenesis^28^. Intriguingly, MuSC activation coincides with neutrophil abundance reaching its peak, suggesting that the neutrophils may be involved in MuSC activation^29^. However, to date, there is no direct evidence to support this hypothesis.

Numerous studies have shown that macrophages or neutrophils regulate the dynamics of MuSCs via paracrine factor release^28,30,31,32,33,34,35^. However, only a few studies have reported that cell-cell contacts play an essential role in regulating the fate of MuSCs. For example, it has been shown that macrophages protect human myogenic cells from apoptosis through direct cell-cell communications *in vitro*^36,37^. In addition, *in situ* ultrastructural analysis indicated that skeletal muscle regeneration involved cell-to-cell interaction between activated macrophages and satellite cells/myogenic precursor cells in the proliferating and differentiating stage^38^. Furthermore, the adhesion molecule VCAM-1 and its highest-affinity ligand VLA-4 mediated interaction between MuSCs and CD45^+^ immune cells, mainly neutrophils and macrophages, which may play an essential role in muscle regeneration^39,40^. However, these studies of the role of macrophages or other myeloid cells during muscle regeneration were mainly based on *in vitro* or *ex vivo* analysis and didn’t involve the early activation stage of MuSCs. A recent breakthrough was made by using real-time imaging of muscle injury models in transgenic zebrafish to systematically capture the interactions between MuSCs and the innate immune system. D. Ratnayake *et al*. reported that all MuSCs at wound sites must establish prolonged, direct interaction with a specific resident macrophage before proper cell division and the contact ceased upon cell division after injury in larval zebrafish^41^. By contrast, the origins of mammals’ skeletal muscle resident macrophages (SMRMs) are heterogeneous^42,43^, it is extremely challenging to find a transgenic model that labels all SMRMs, infiltrating macrophages, and neutrophils simultaneously. Therefore, the spatiotemporal interaction between myeloid cells and MuSCs in mammals has not been investigated *in vivo* in real-time, and it is unknown whether constant macrophage or other myeloid cell-MuSC contact is also present and, if so, necessary for MuSC activation and proliferation in response to muscle injury in mammals.

In this study, we developed a multimodal non-linear optical microscope system integrating second harmonic generation (SHG) and two-photon excited fluorescence (TPEF) imaging to examine the dynamics of satellite cells, non-myogenic cells and their complex interactions during muscle regeneration *in vivo*. Combining signals from fluorescent proteins of transgenic mouse models and the reduced nicotinamide adenine dinucleotide (NADH), a metabolic coenzyme located in the mitochondria with autofluorescence, in principle, we could trace all the cell types that interacted with MuSCs/activated satellite cells (ASCs) in the injured sites. As a result, we first monitored the change in mitochondrial mass and volume of MuSCs in vivo and found they increased synchronously after injury. Further, we demonstrated that cell-cell contacts between RFP^+^ monocytes/macrophages and MuSCs were not essential for the cell growth of MuSCs during early activation. Notably, we found that MuSC proliferation in mice did not require continuous contact with a specific macrophage or other non-myogenic cells as in zebrafish. However, depletion of macrophages significantly impaired the proliferation and differentiation of MuSCs and led to intramuscular fibrosis. Besides, by reducing the infiltration of neutrophils in the CCR2 knockout (ko) mice, we demonstrated that the infiltrating neutrophils and macrophages had no influence on the initial cell volume increase in MuSCs. These new observations reveal the stage-dependent role of MuSC-myeloid cell crosstalk during muscle regeneration.

## Results

### *In vivo* dynamics of MuSCs during skeletal muscle regeneration

To study the dynamics of MuSCs during muscle regeneration *in vivo*, we used a multimodal two-photon microscope system to visualize genetically labeled Pax7^+^ MuSCs and their descendant myoblasts in live mice (Supplementary Fig. 1). Pax7^+^ SCs were efficiently labeled by cytoplasmic eYFP after tamoxifen injection^44,45^. Additionally, reduced nicotinamide adenine dinucleotide (NADH) is much more abundant in mitochondria than in other organelles. Therefore, its autofluorescence can be used as a label-free “mitotracker” probe for visualization of the distribution of mitochondria in cells^46,47,48^. The second harmonic generation (SHG) signals from muscle sarcomeres and collagen fibers were used to visualize the regeneration process^49,50,51^.

We performed two-photon imaging on mouse hind limb tibialis anterior (TA) muscle at different time points after needle track injury. Before injury, quiescent MuSCs resided on the surface of individual muscle fibers, indicated by the orderly arranged SHG pattern from sarcomeres (Fig. 1a). 12-hour post needle track injury (12 hpi), the muscle fiber degenerated, and the SHG signal from sarcomeres disappeared. However, we observed that some collagen fibers parallel to the adjacent intact myofiber were still present in the injured site, as shown by arrows in Fig. 1a. These collagen fibers were referred to as “ghost fibers”, the longitudinal axis of which determined the direction of migration and division of MuSCs after injury^44^. Previously, based on freshly isolated MuSCs or myofibers grown in culture, activated MuSCs were found to be larger in volume than their quiescent counterparts with expanded cytoplasm and more organelles^45,52,53,54,55,56^. Here, we monitored the cell volume change in live mice to investigate the kinetics of MuSCs activation and growth in response to acute injury *in vivo*. At 12 hpi, a slight increase in cell volume compared to uninjured cells marked one of the earliest visible signs of MuSC activation (the mean value increased by about 29%). Then, the cell volume increased continuously, reaching its maximum at 2 dpi when MuSCs started to proliferate, and subsequently decreased, returning to pre-injury levels at 1 mpi (Fig. 1b). By analyzing the autofluorescence of NADH in individual YFP^+^ cells after injury, we also observed that the increased cell volume was accompanied by increased mitochondrial mass (Fig. 1c and d). The distribution of nuclei in myocytes and newly formed myotubes could also be visualized (Fig. 1d, indicated by the asterisks). Our *in vivo* data showed that proliferating myoblasts and differentiating myocytes underwent enhanced mitochondrial biogenesis, consistent with previous *in vitro* analyses^45,57^.

**Fig. 1:**
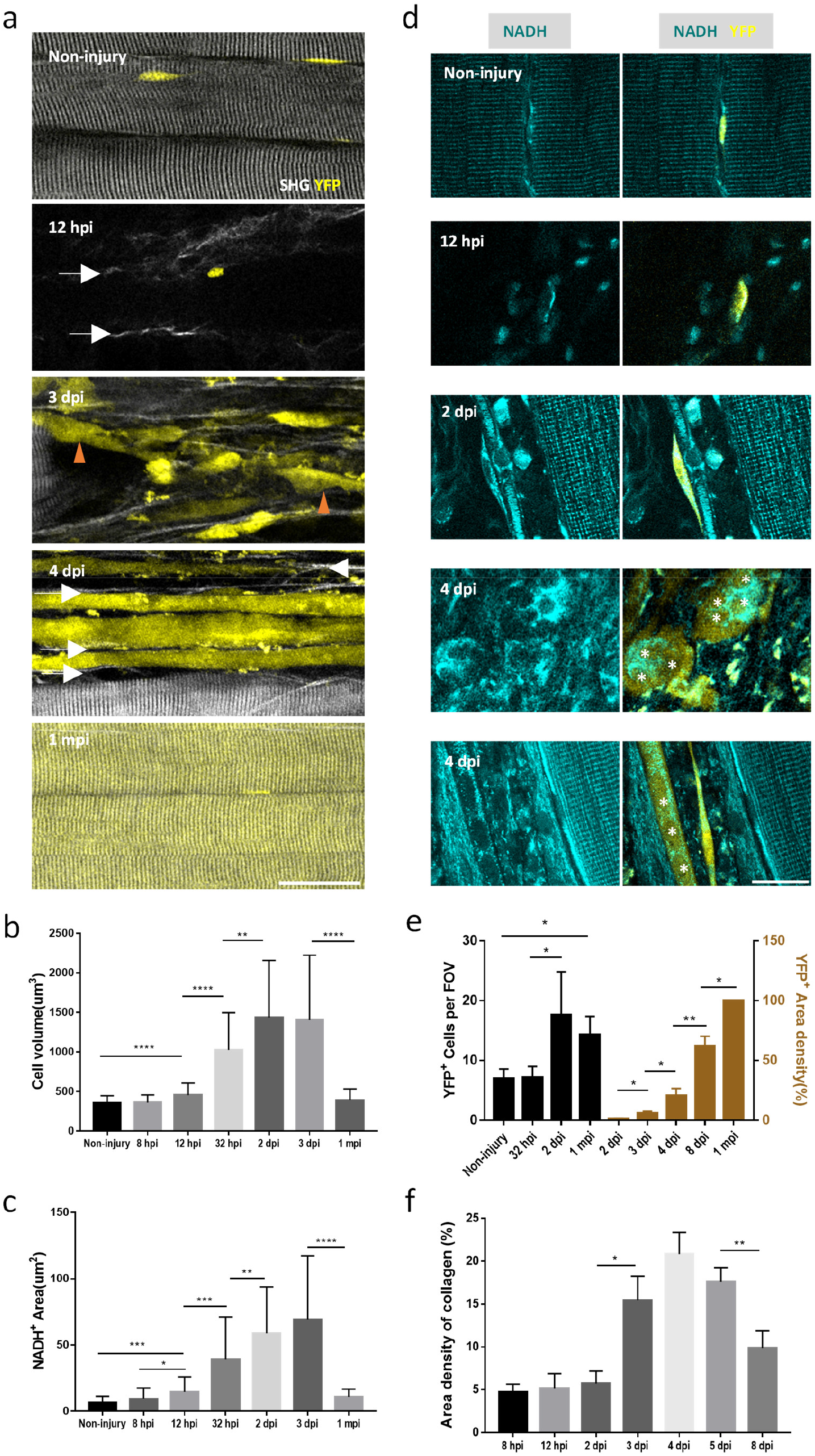
The dynamic of MuSCs and their microenvironment during muscle regeneration. (a) Typical YFP (yellow) and SHG (grey) images of muscle satellite cells/myogenic progenitor cells/myotubes and muscle environment at indicated time points after injury. Collagen parallel to the intact muscle fiber was indicated by arrows. Myoblast that fused together indicated by brown arrow heads. (hpi and dpi, hours/days post injury.) (b) Time course of cell volume of individual YFP^+^ at indicated time points; ≥30 cells from 3 mice per time point. (c) Time course of NADH area (mitochondrial mass) of individual YFP^+^ muscle satellite cells/myogenic progenitor cells at indicated time points. ≥30 cells from 3-5 mice per time point. (d) The NADH signal (cyan) of representative YFP^+^ muscle satellite cells/myogenic progenitor cells/myotubes (yellow). Asterisks show the nuclei distribution in the myocytes/myotubes at 4 dpi. (e) Time course of YFP^+^ cell density (left y-axis, black) and YFP^+^ area density (right y-axis, brown) at different time points. (n ≥3 mice per time point) (f) The area density of collagen of the injured sites at indicated time points. (n ≥3 mice per time point). The FOV is 300 μm*300 μm*60 μm. Data were reported as mean ± SD (Standard deviation). (*: p<0.05; **: p<0.01; ***: p<0.001; ****: p<0.0001; student t-test). Scale bars: 50 μm in (a), 30 μm in (d).

To evaluate proliferation and differentiation of MuSCs, we measured the cell density of individual YFP^+^ cells before 2 dpi and calculated the percentage of YFP^+^ area of the injured sites after 2 dpi (Fig. 1e). Cell density at 32 hpi was similar to that of MuSCs in non-injured muscle and increased slightly at 2 dpi. At 3 dpi, YFP^+^ cell density increased significantly, and some cells started to fuse (Fig. 1a). After 4 dpi, newly formed myotubes were surrounded by collagen fibers parallel to the intact muscle cells (Fig. 1a, Supplementary Fig. 2a). By analyzing the SHG signal generated by type I and III collagen^58^ (Supplementary Fig. 2b), we found that the collagen density in the injured sites increased significantly at 3 dpi, remaining high until 5 dpi and decreased at 8 dpi (Fig. 1f). The increase in collagen content may promote myoblast migration and myogenic differentiation^59^. At 1 mpi, YFP^+^ muscle fibers filled the injured sites, and the periodic SHG signal indicated that the sarcomeres were regenerated (Fig. 1a). The non-myofiber YFP^+^ cell (i.e., self-renewed MuSCs) density was slightly higher at 1 mpi than that at the non-injury stage (Fig. 1e). In addition, based on the polarization dependence of SHG intensity^60^, we enhanced the SHG intensity of collagen fiber and weakened the SHG intensity of the muscle fibers by changing the polarization of the excitation laser. Then we could quantify the collagen density before injury and 1 mpi and found no significant difference between them (Supplementary Figs. 2c-d). In summary, our multimodal two-photon microscope is an excellent tool for studying the dynamics of MuSCs and the environment of the muscle system during muscle regeneration *in vivo*.

### The interaction between RFP^+^ monocytes/macrophages and MuSCs during MuSC activation

After an acute injury, various immune cells, predominantly neutrophils and monocytes/macrophages (MOs/MΦs), infiltrate the injured muscle. The neutrophils invade the injured muscle first and almost disappear before 2 dpi^8,9,10^. On the other hand, monocytes/macrophages that express pro-inflammatory markers infiltrate injured muscle fibers via the CCL2-CCR2 axis^12,14,15^. Gradually, these pro-inflammatory macrophages were outnumbered by the anti-inflammatory macrophages by 7 dpi. Therefore, to explore the role of monocytes/macrophages and the interaction between them and MuSCs, we performed *in vivo* imaging in a transgenic mouse model (Pax7^CreERT2/+:^ Rosa-stop-YFP; Ccr2^RFP/+^) in which the CCR2^+^ MOs/MΦs were labeled by RFP^61^, and Pax7^+^ MuSCs were marked by YFP after tamoxifen injection.

It was reported that macrophages could release ADAMTS1 that targets Notch1 protein of the MuSCs to affect the activation of MuSCs^32^. In addition, MOs/MΦs can also establish cell-cell connections with MuSCs through membrane ligands and receptors, which may regulate the lineage progression of MuSCs after injury^39,62^. However, whether direct cell-cell contact between MuSCs and monocytes/macrophages is required for the activation of MuSCs is uncertain.

As mentioned previously, the cell volume of MuSCs increased significantly at 12 hpi but not at 8 hpi. Therefore, we performed time-lapse imaging in the TA muscle from 4.5 hpi to 12 hpi to monitor the dynamics of RFP^+^ MOs/MΦs, the spatiotemporal interaction between MuSCs and RFP^+^ MOs/MΦs and to trace the volume of individual MuSCs. However, we failed to observe the recruitment of RFP^+^ MOs/MΦs when we only kept the body temperature of mice by using a heating pad set to 36°C (Supplementary Fig. 3a). The low temperature of the hind limb (~23°C) and high pressure on the muscle impaired the blood flow, further impeding the immune response^63,64^. Therefore, we redesigned a mount to keep the temperature of both the body and the exposed TA muscle at normal values (35-36°C and 28-30°C, respectively) and to reduce the pressure of the coverslip on the muscles (Supplementary Figs. 3b-e). Finally, we could monitor the *in vivo* dynamics of cells for up to 7.5 hrs.

As the results showed, there were few RFP^+^ MOs/MΦs resident in the intramuscular connective tissue before injury (Supplementary Fig. 4a). The density of RFP^+^ MOs/MΦs slightly increased at 4.5 hpi compared to the non-injury state (Supplementary Fig. 4b). From 4.5 to 12 hpi, the MuSCs were motionless (Supplementary Fig. 4c), and RFP^+^ MOs/MΦs were gradually recruited into the injured sites in Injured-Ctrl groups (Fig. 2a-b). Most RFP^+^ MOs/MΦs were highly dynamic (Supplementary Fig. 4d, Supplementary Video 1). Moreover, a small population of RFP^+^ MOs/MΦs remained in the injured site during imaging, and the displacement was less than 10 μm (Supplementary Fig. 4e, Supplementary Video 2). No division of RFP^+^ MOs/MΦs was observed during imaging, consistent with the *ex vivo* results that there were no Ki67^+^ monocytes/macrophages before 15 hpi^14^. We recorded the cell volume of individual MuSCs at 4.5 hpi, 8 hpi and 12 hpi, and analyzed the minimum distance between MuSCs and RFP^+^ MOs/MΦs every 5 minutes from 4.5 to 12 hpi (Fig. 2c-d). When MuSCs were in contact with RFP^+^ MOs/MΦs, the minimum distance was zero. However, it should be emphasized that direct contact between MuSCs and macrophages cannot be confirmed definitively due to optical microscopy’s limited resolution. By analyzing the change in distance between them, we found that all MuSCs established short-lived rather than continuous or repeated direct interaction with RFP^+^ MOs/MΦs before cell volume increased at 12 hpi (Fig. 2c-d, Supplementary Video 3). In addition, MuSCs interacted directly with different RFP^+^ MOs/MΦs but only one or two at a time immediately after injury (Supplementary Fig. 4f). We found that the increase in cell volume of different MuSCs was asynchronous, which may be caused by the unsynchronized muscle degeneration and intrinsic heterogeneity of MuSCs^55,56^. To exclude the possibility that long-term *in vivo* imaging affected MuSC activation and altered its interaction with macrophages, we conducted time-lapse imaging on non-injured muscles (Supplementary Fig. 4g, Supplementary Video 4). In intact muscle fibers, the cell volume of MuSCs did not increase significantly compared to the Injured-Ctrl group after 7.5-hour imaging (Fig. 2c), and a portion (5 of 12 cells in 3 mice) of MuSCs was observed to contact different RFP^+^ MOs/MΦs during the imaging session (Fig. 2d). In addition, the duration of contact between MuSCs and RFP^+^ cells was similar to that in injured muscle (Supplementary Figs. 4h-i). Together, MuSCs did establish short-lived direct contacts with RFP^+^ cells during activation immediately after injury.

**Fig. 2:**
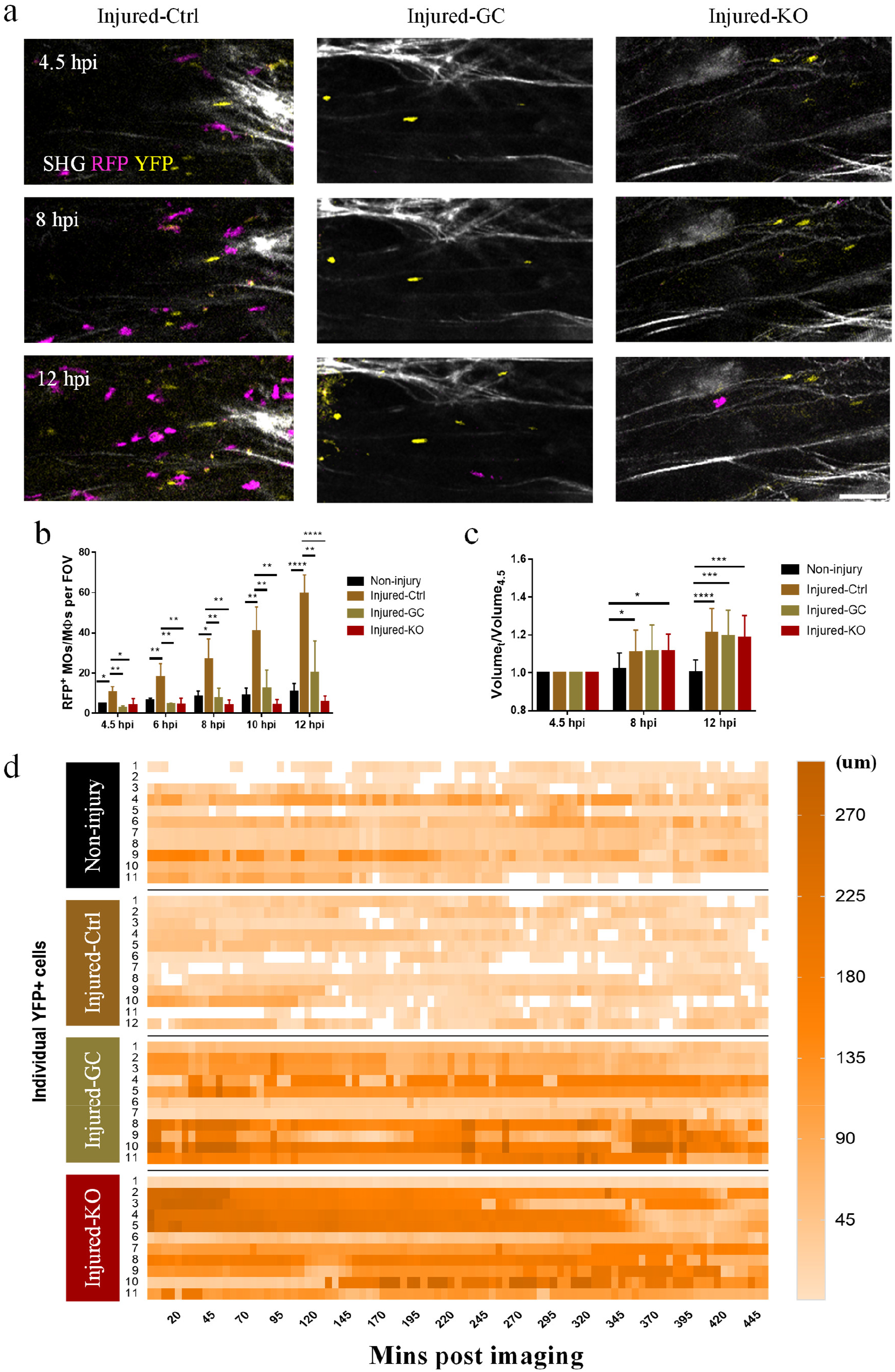
The spatiotemporal interaction of RFP^+^ monocytes/macrophages (MOs/MΦs) and MuSCs immediately after injury (from 4.5 to 12 hpi). (a) Maximum z intensity projections of TPEF image stacks of YFP^+^ MuSCs (yellow), RFP^+^ MOs/MΦs (magenta) and SHG generated from collagen and muscle fiber (grey) at 4.5 hpi, 8 hpi and 12 hpi in Injured-Ctrl, Injured-GC (Glucocorticoid), and Injured-KO (CCR2 knockout) groups. The image started at 4.5 hpi. (b) The time course of RFP^+^ MOs/MΦs density at the image sites during time-lapse imaging in the Non-injury, Injured-Ctrl, Injured-GC, and Injured-KO groups. n=3 mice per groups. The FOV is 300 μm*300 μm*60 μm. (c) The change in normalized cell volume of individual MuSCs during time-lapse image in four groups. (n≥12 cells from 3 mice per groups) (d) The time course of minimal distance from representative individual MuSCs to RFP^+^ MOs/MΦs during time-lapse imaging in Non-injury, Injured-Ctrl, Injured-GC, and Injured-KO groups. The minimal distances with a value of 0 um were highlighted with white squares. Other values were coded with color maps. The volume of the 3D distance matrix equals to the FOV. Data were reported as mean ± SD. (*: p<0.05; **: p<0.01; ***: p<0.001; ****: p<0.0001; student t-test). Scale bars: 50 μm in (a), 30 μm in (b).

To further explore whether the direct cell-cell contacts between RFP^+^ MOs/MΦs and MuSCs were required to activate MuSCs, we tried to partially reduce macrophage infiltration by glucocorticoid (GC) treatment (Injured-GC group)^24,65^ or using CCR2-deficient (CCR2ko) mice (Injured-KO group)^20^. We repeated time-lapse imaging in the Injured-GC and Injured-KO mice. The infiltration of RFP^+^ MOs/MΦs during imaging was significantly impaired in the Injured-GC and Injured-KO groups as only a few RFP^+^ cells were recruited to the injured areas at 12 hpi (Fig. 2a-b, Supplementary Video 5 and 6). Further, because of the low density of RFP^+^ MOs/MΦs, most MuSCs could not contact RFP^+^ MOs/MΦs directly before 12 hpi (Fig. 2d, Supplementary Video 7 and 8). However, there was no difference in the increase in cell volume between MuSCs of the Injured-Ctrl group that had been in direct contact with RFP^+^ MOs/MΦs and MuSCs of the Injured-GC and Injured-KO group that had not directly contacted RFP^+^ MOs/MΦs before 12 hpi (Fig. 2c). (There were about ~2/15 and ~1/13 MuSCs that contacted RFP^+^ MOs/MΦs before 12 hpi in the Injured-GC or Injured-KO groups respectively, and these MuSCs were excluded.) To investigate whether the reduction in infiltrated macrophages impaired cell growth in later stages, we performed snap-shot imaging at 0 hpi, 8 hpi, 12 hpi, 2 dpi and 3 dpi and analyzed MuSC volume. The RFP^+^ cell density was significantly reduced in the Injured-GC and Injured-KO groups before 12 hpi (Supplementary Fig. 4j). Except that the volume of MuSC in the Injured-GC group was slightly less than that of the Injured-Ctrl group at 2 dpi (mean value decreased by ~12%), MuSC growth was similar in all these groups at all time points examined (Supplementary Fig. 4k). Collectively, our data indicated that both the decrease in RFP^+^ MOs/MΦs density and the loss of direct cell-cell contact between MuSCs and RFP^+^ MOs/MΦs did not impede the activation of MuSCs. Besides, the reduction in recruited macrophages did not impair the cell growth of MuSCs.

### The interaction between RFP^-^NADH^+^ non-myogenic cells and MuSCs during MuSC activation

During time-lapse imaging sessions, along with RFP^+^ MOs/MΦs, a significant number of RFP^-^NADH^+^ YFP^-^ cells was observed to infiltrate the injured sites in the Injured-Ctrl, Injured-GC and Injured-KO groups (Fig. 3a). Their density was higher than RFP^+^ MOs/MΦs at 4.5 hpi. To confirm that the NADH signal could be used to visualize most cells in the injured muscles, Mito Tracker Deep Red (MTDR) that could stain mitochondria in live cells, was applied topically to stain cells *in situ*. The results showed that more than 90% of cells were NADH^+^MTDR^+^ (Supplementary Figs. 5a-b). Therefore, most cells could be visualized based on NADH autofluorescence without further labeling. Since the NADH signal was mainly generated by mitochondria inside cells, it could not delineate the entire cell boundary. By analyzing the RFP/YFP labeled whole cells and their NADH signal, we found the difference between the boundary of NADH signal and RFP/YFP signal was probably less than 2.5 μm laterally and 6 μm axially (Upper 95% confidential interval of mean value were 2.165 μm and 3.101 μm), respectively (Fig. 1b and Supplementary Figs. 5c-d). Therefore, if the distance between NADH^+^ cells and MuSCs was greater than 2.5 μm laterally and 6 μm axially, there was no direct contact between cells.

**Fig. 3:**
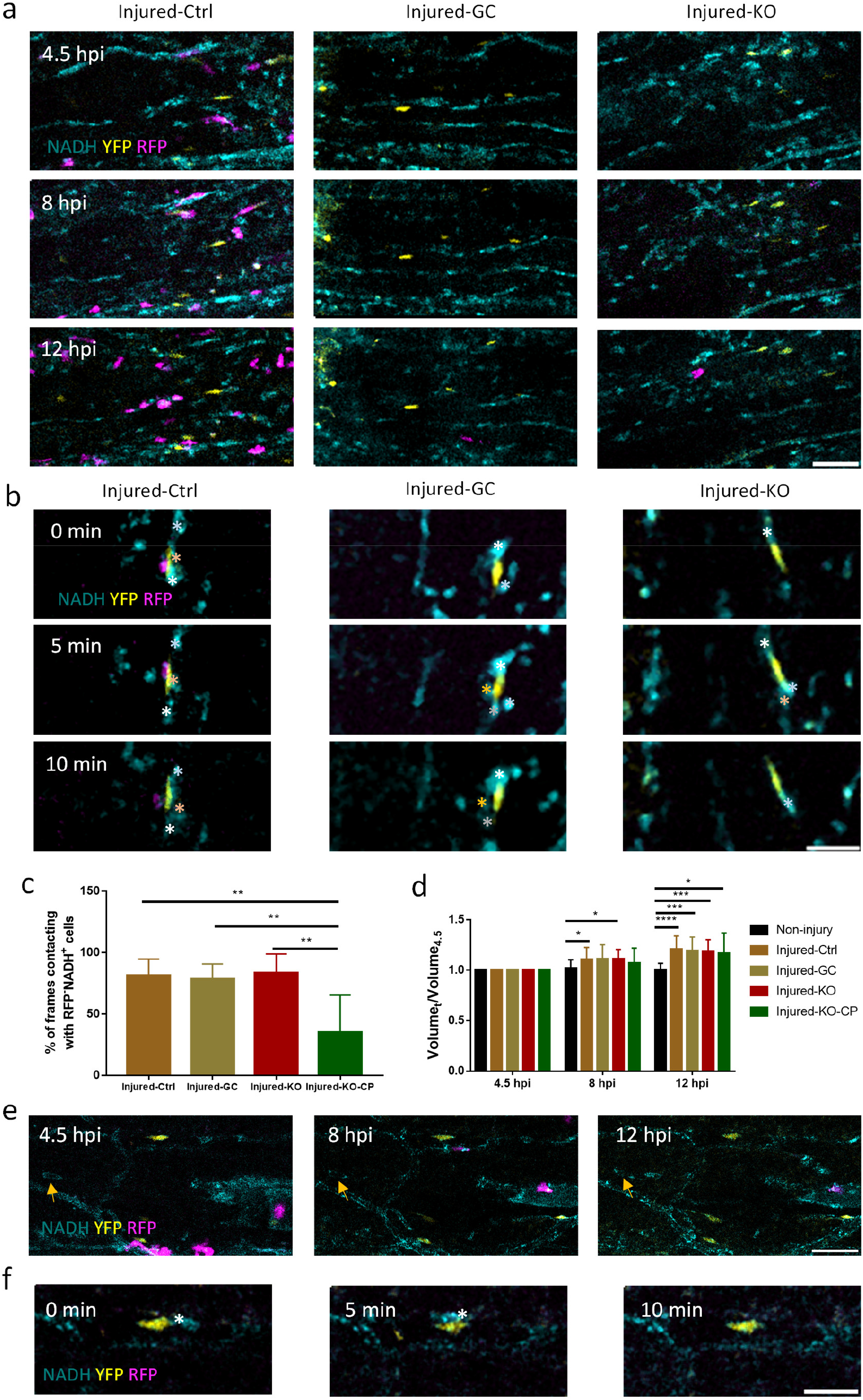
The spatiotemporal interaction of RFP^-^NADH^+^ non-myogenic cells and MuSCs immediately after injury. (a) Maximum z intensity projections of TPEF image stacks of YFP^+^ MuSCs (yellow), RFP^+^ MOs/MΦs(magenta) and NADH^+^ cells (cyan) at 4.5 hpi, 8 hpi and 12 hpi in Injured-Ctrl, Injured-GC, and Injured-KO groups. (b)Typical images for the interaction between RFP^-^NADH^+^ cells and MuSCs within 10 mins in the three group. Different color asterisks corresponded to different non-myogenic cells at different time points. Cyan, NADH; yellow, YFP; magenta, RFP. (c) The total contact duration between RFP^-^NADH^+^ cells and MuSCs in the Injured-Ctrl, Injured-GC, Injured-KO and Injured-KO-CP (CCR2ko mice treated with Cyclophosphamide (CP)) groups. (d) The change in normalized cell volume of individual MuSCs during time-lapse image in the Non-injury, Injured-Ctrl, Injured-GC, Injured-KO and Injured-KO-CP groups. (n ≥ 12 cells from 3 mice per groups). No significant difference was observed between these groups. (e) Maximum z intensity projections of TPEF image stacks of YFP^+^ MuSCs (yellow), RFP^+^ macrophages (magenta) and NADH^+^ cells (cyan) at indicate time post injury in the Injured-KO-CP groups. The brown arrows indicated the motionless RFP^-^NADH^+^ non-myogenic cell. (f) TPEF images for the interaction between RFP^-^NADH^+^ cells and MuSCs within 10mins in the Injured-KO-CP groups. Asterisks indicated an RFP^-^NADH^+^ cell that contacted the MuSCs within 10 minutes. Data were reported as mean ± SD. (*: p<0.05; **: p<0.01; ***: p<0.001; ****: p<0.0001; student t-test). Scale bars: 50 μm in (a) (e), 30 μm in (b), (f).

As demonstrated in previous studies, neutrophils were the earliest immune cells to infiltrate skeletal muscle in response to injury^8^. Therefore, we immunostained the injured TA muscle using the markers Gr-1 and Ly6G, respectively. More than 83% of these cells were found to be Ly6G^+^ or Gr-1^+^ neutrophils (Supplementary Figs. 5e-f). Due to the high density and dynamic nature of RFP^-^NADH^+^ YFP^-^ cells, they started to interact with MuSCs frequently at 4.5 hpi, and occasionally, MuSCs were observed to contact multiple cells simultaneously (Fig. 3b and Supplementary Videos 3, 7 and 8). Further, there was no significant difference in the total duration of direct contacts between MuSCs and NADH^+^RFP^-^ cells among Injured-Ctrl, Injured-GC and Injured-KO groups (Fig. 3c).

To further examine whether the interaction between MuSCs and RFP^-^NADH^+^ cells was essential for MuSCs activation, we tried to deplete neutrophils in the CCR2ko mice. A standard *in vivo* method of neutrophil depletion is anti-Ly6G antibody-mediated depletion^66,67^. But we failed to deplete the neutrophils by using clone 1A8 in the CCR2ko mice (data not shown) since macrophages are required for effective anti-Ly6G-mediated depletion of neutrophils^68^. Therefore, we tried another drug, Cyclophosphamide (CP), which can trigger the formation of DNA crosslinks and lesions that induce cell death. Injection of CP could also reduce the number of circulating monocytes, B and T cells by limiting the proliferation of dividing cells^69,70^. However, the density of B and T cells was much lower than neutrophils and macrophages during the early stage of injury-induced muscle regeneration as B and T cells are recruited to the injured site after 3 dpi^9,10,11,71^. Therefore, the side effects of CP on B and T cells may not be of concern before 12 hpi. Treatment of CP effectively reduced the number of RFP^-^ NADH^+^ cells (neutrophils) at 4.5 and 12 hpi in the Injured-KO group (Supplementary Fig. 5g). We next repeated the time-lapse imaging in the CP-treated CCR2ko (Injured-KO-CP) mice and found that the cell volume of MuSCs in the Injured-KO-CP groups was not significantly different from that in the Injured-Ctrl, Injured-GC and Injured-KO groups (Fig. 3d), although both the density of RFP^-^NADH^+^ cells and the frequency of contact were reduced by comparison with the other three groups (Fig. 3c, e-f, Supplementary Videos 9 and 10). Therefore, the reduction of neutrophil density and contact duration between MuSCs and RFP^-^NADH^+^ cells had no influence on the initial growth of MuSCs after injury. In addition, we observed that some RFP^-^NADH^+^ cells were motionless as MuSCs (Fig. 3e), and they had a similar pattern to MuSCs in the NADH channel (Supplementary Figs. 5h-i). Besides, some motionless non-myogenic cells established constant contact with MuSCs (Supplementary Fig. 5j). It is unclear whether this contact had any effect on MuSC activation.

### The interaction between RFP^+^/RFP^-^ non-myogenic cells and activated satellite cells (ASCs) during the proliferation stage of muscle regeneration

The reduction of infiltrated monocytes/macrophages and the interaction between them with MuSCs did not impair the initial volume increase and the further growth of MuSC. Therefore, we tried to determine whether proliferation of MuSCs would be impeded.

Infiltration of macrophages preceded the proliferation of ASCs. At 1 dpi, many RFP^+^ monocytes/macrophages were recruited to the injured site, and granules with strong auto-fluorescence in the YFP channel were located inside some RFP^+^ MOs/MΦs (Fig. 4a and Supplementary Fig. 6a). The autofluorescence may be generated from NAD(P)H, FAD and lipofuscin inside the macrophages’ cytoplasm after the phagocytosis of muscle fiber debris^72,73,74^. More RFP^+^ MOs/MΦs infiltrated the injured areas and surrounded the YFP^+^ activated satellite cells (ASCs) at 1.5 dpi and 2 dpi (Fig. 4a). Besides, most NADH^+^ areas were RFP^+^ after 1 dpi in the Injured-Ctrl group, suggesting that monocytes/macrophages were the predominant cell type (Supplementary Fig. 6b). ASC density increased significantly at 2-3 dpi, indicating that cell divisions were relatively frequent starting at 2 dpi. Thus, we first conducted time-lapse imaging to monitor the spatiotemporal interaction between macrophages and ASCs at 2 dpi every 5 minutes for 7.5 hrs. Supplementary Fig. 6c and Supplementary Video 11a showed that a single ASC migrated parallel to intact neighboring muscle fibers and was continuously surrounded by macrophages before division. Unfortunately, due to the high density of RFP^+^ MOs/MΦs, it was hard to identify individual macrophages to study their interaction with ASCs. There was a special case in which the muscle fiber degenerated during imaging, and an ASC emerged without being surrounded by many RFP^+^ MOs/MΦs like other ASCs did at 2 dpi. During the time-lapse imaging, this ASC only migrated within a short distance, and it seemed that a small cluster of RFP^+^ MOs/MΦs below the ASC contacted it repetitively before cell division (Supplementary Fig. 6d, Supplementary Video 11b). However, it was hard to determine whether there was a specific RFP^+^ cell contacting ASC constantly due to the higher density of RFP^+^ MOs/MΦs in the late imaging period.

**Fig. 4:**
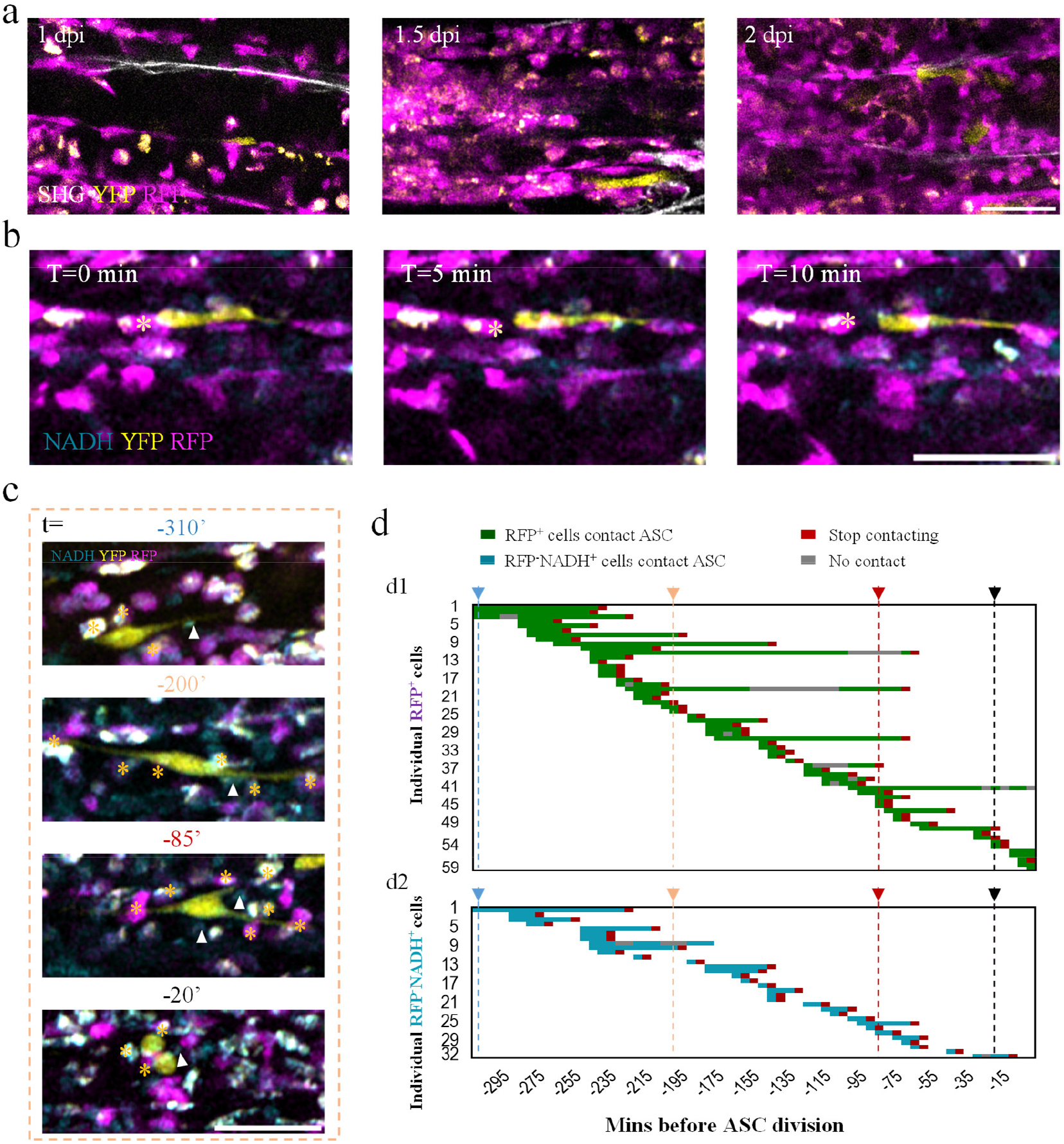
The spatiotemporal interaction between RFP^+^/RFP^-^ non-myogenic cells and YFP^+^ activated satellite cells (ASCs) at the proliferation stage in the Injured-Ctrl groups. (a) Representative images indicated the infiltration of macrophages at 1dpi, 1.5 dpi and 2 dpi. (RFP, magenta; YFP, green; SHG, grey; autofluorescence within macrophages, white). (b) Representative time sequence images of the dynamic contact between non-myogenic cells and MPs at 1 dpi. (RFP, magenta; YFP, yellow; NADH, cyan) Asterisks indicated a RFP^+^ MO/MΦ that established a short-lived contact with ASC at 1 dpi. (c) Image sequence of the interaction between a traced ASC and non-myogenic cells in the control group at 1.5 dpi. Asterisks indicated the RFP^+^ MOs/MΦs that contacted the ASC at the indicated time points. White arrow heads indicated the RFP^-^NADH^+^ cells that contacted the ASC at the indicated time points. Daughter cells separated from each other at time 0. (d) Representative compilation of the spatiotemporal interaction between different RFP^+^ MOs/MΦs (d1), RFP^-^NADH^+^ cells (d2) and the traced ASC shown in (c). The different color arrows and dotted lines correspond to the different time points shown in the (c). Each row represented an RFP^+^ MO/MΦ (d1) or an RFP^-^NADH^+^ cell (d2). Green boxes or cyan boxes mean that the RFP^+^ MOs/MΦs or RFP^-^NADH^+^ cell establishes contact with the ASC and red box means the contact is stopped before this frame. The number of the green or cyan boxes that the dotted lines pass through equal to the number of RFP^+^ MOs/MΦs or RFP^-^NADH^+^ cells that contacted ASC at the same time. For example, at t=-200’, there were 7 RFP^+^ MOs/MΦs and 1 RFP^-^NADH^+^ cell that contacted ASC. Besides, grey box means that the RFP^+^ MOs/MΦs temporarily dissociated from ASCs at that time but would contact ASCs again. White area means that the non-myogenic cells are not traced. Scale bars: 50 um in (a-b), (e), 30 um in (c).

Importantly, the density of YFP^+^ ASCs at 2 dpi was almost twice of that in non-injured mice, indicating that the first cell division already occurred in most MuSCs by 2 dpi (Fig. 1e). Therefore, we tried to perform the time-lapse imaging at 1-1.5 dpi to better examine the spatiotemporal interaction between RFP^+^ MOs/MΦs and ASCs. However, during 7.5-hour time-lapse imaging at 1 dpi, we did not observe the division of ASCs which were mobile and contacted with different RFP^+^ MOs/MΦs transiently (Fig. 4b and Supplementary Video 12a). After 1.5 dpi, we could observe cell division, but ASCs were surrounded by RFP^+^ MOs/MΦs later as they were at 2 dpi (Supplementary Video 12b). Therefore, we reduced the number of the punctured sites from 5 to 3 on the TA muscles and avoided bleeding to reduce inflammation. Finally, in some cases, we could observe the ASC divisions at 1.5 dpi without being surrounded by RFP^+^ MOs/MΦs. Because the infiltrating macrophages were the dominant cell type in the injured sites, we first analyzed the dynamics of individual RFP^+^ MOs/MΦs and the ASCs before ASC division. Due to the high density of MOs/MΦs, a traced ASC contacted different RFP^+^ MOs/MΦs simultaneously at different time points (Fig. 4c, d1 and Supplementary Video 12c). The compilations of the behavior of 59 RFP^+^ MOs/MΦs that contacted the traced ASC before ASC division were shown in the Fig. 4d1. At t = -200’ (Daughter cells separated with each other at t = 0’), the ASC contacted seven RFP^+^ MOs/MΦs simultaneously. No RFP^+^ MOs/MΦs established constant contact with the ASC before cell division, and the contact duration was lower than 3 hours (Supplementary Fig. 6e).

As mentioned before, the recruited monocyte-derived macrophages were CCR2^+^ and RFP^+^, but most resident macrophages were CCR2^- 42,75^. Therefore, to study the interaction between all macrophages and ASCs, we also need to monitor the dynamics of RFP^-^ macrophages. As mentioned before, more than 90% NADH^+^ cells were RFP^+^ after 1 dpi. There were still some RFP^-^NADH^+^ cells in the injured sites. And due to the long-term exposure of the muscle, more RFP^-^NADH^+^ cells (likely neutrophils) migrated to the injured sites (Supplementary Fig. 6f). Thus, we also analyzed the interaction between RFP^-^ NADH^+^ cells and ASCs. It should be noted that if the RFP^-^ non-myogenic cells had long protrusion as ASCs (as shown in t=-200’ in Fig. 4c), and the protrusions could not entirely be observed by NADH signal, the contact would be missed. Similar to RFP^+^ MOs/MΦs, RFP^-^NADH^+^ cells were highly dynamic and established transient contact with the ASCs (Fig. 4c and d2). Hereinto, the contact duration between RFP^-^NADH^+^ cells and the ASCs was significantly shorter than that between RFP^+^ MOs/MΦs and the ASCs (Supplementary Fig. 6e). Therefore, our data strongly suggest that the constant contact between ASCs and non-myogenic cells is not required for the ASCs division.

To better address the role of macrophages in ASCs division, we reduced macrophage infiltration by glucocorticoid (GC) treatment for several days and using *CCR2ko* mice. We found that RFP^+^ MOs/MΦs were significantly reduced in either Injured-GC or Injured-KO mice at 2 dpi (Fig. 5a-b). The density of ASCs in macrophage-depleted mice was slightly lower at 2 dpi compared with Injured-Ctrl mice but still higher than that in non-injured mice (Fig. 5c). To study potential interaction between ASCs and individual infiltrating macrophages before ASCs underwent the first cell division, we conducted time-lapse imaging at 1.5 dpi in Injured-GC and Injured-KO mice, analyzing the dynamics of macrophages and ASCs from when ASCs were observed to when ASCs completed cell division (Fig. 5d-i). Before cell division, ASCs only established short-lived contact with several RFP^+^ MOs/MΦs or did not contact RFP^+^ MOs/MΦs during imaging in Injured-KO mice because of their lower cell density (Fig. 5d and g1, Supplementary Fig. 6g, Supplementary Video 13). The maximum duration that RFP^+^ MOs/MΦs contacted ASCs was less than 40 mins in the Injured-KO mice (Supplementary Fig. 6e).

**Fig. 5:**
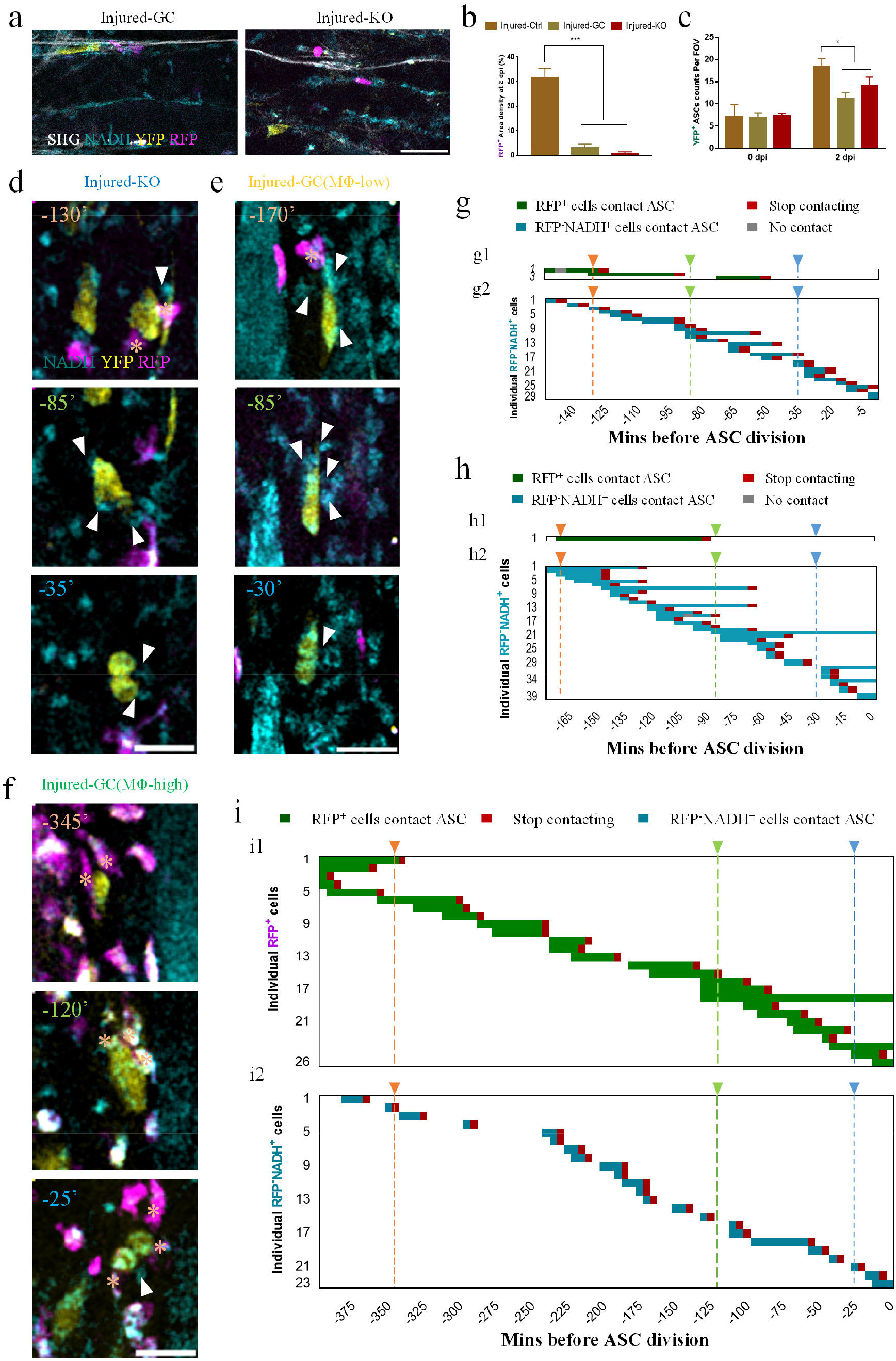
The spatiotemporal interaction between RFP^+^ /RFP^-^non-myogenic cells and YFP^+^ activated satellite cells (ASCs) at the proliferation stage in the Injured-GC and Injured-KO groups. (a) Representative image for the TA muscle at 2 dpi of Injured-GC and Injured-KO groups which indicated the successful depletion of macrophages. SHG, grey; YFP, yellow; NADH, cyan; RFP, magenta. (b) RFP^+^ MOs/MΦs density of three groups. (n ≥3 mice per time point). (c) YFP^+^ ASCs density at non-injury and 2 dpi of three groups. (n ≥3 mice per time point). (d-f) Representative time sequence images of the dynamic contact between non-myogenic cells and ASCs before myoblast division at 1.5 dpi in Injured-KO (d) Injured-GC groups with low (e) and high (f) RFP+ MΦ density.. Daughter cells separated from each other at time 0. (RFP^+^ MOs/MΦs, magenta; YFP^+^ ASCs, yellow; NADH, cyan). Asterisks indicated the RFP^+^ MOs/MΦs that contacted ASC at the indicated time points. White arrow heads indicated the RFP^-^NADH^+^ cells that contacted ASC at the indicated time points. (g-i) Representative compilation of the spatiotemporal interaction between non-myogenic cells & ASCs before cell division in the Injured-KO (g) and Injured-GC groups with low (h) and high (i) RFP+ MΦ density at 1.5 dpi. The different color arrows and dotted lines in (g-i) correspond to the different time points in the (d-f) respectively. Daughter cells separated from each other at time 0. Each row represented an RFP^+^ MO/MΦ (g1,h1,i1) or an RFP^-^ NADH^+^ cell (g2,h2,i2). Green boxes or cyan boxes mean that an RFP^+^ MO/MΦ or an RFP^-^NADH^+^ cell establishes contact with the ASC and red box means the contact is stopped before this frame. Similarly, the number of the green or cyan boxes that the dotted lines pass through equal to the number of RFP^+^ MOs/MΦs or RFP^-^NADH^+^ cells that contacted ASCs at the same time. For example, at t=-130’ in the Injured-KO group (d), there were 2 RFP^+^ MOs/MΦs and 1 RFP^-^NADH^+^ cell contacted ASC at the same time (g). Grey box means that the RFP^+^ /RFP^-^ non-myogenic cells do not contact ASCs at that time but contact ASCs again at another time. White area means that the cells are not traced after without contact with ASCs. The different color arrow heads in the (g-i) corresponded to the different time points in the (d-f). Scale bar, 50 um in (a), 30 um in (d-f).

On the other hand, due to the highly variable depletion efficiency by GC treatment, the RFP^+^ cell density fluctuated in Injured-GC mice but remained significantly higher than that in the Injured-KO mice. Therefore, the frequency and duration of contact were higher in the Injured-GC mice (Supplementary Figs. 6e, g). When the RFP^+^ cell density in Injured-GC mice was lower, the ASCs contacted one RFP^+^ cell shortly at one time during imaging (Fig. 5e and h1 and Supplementary Video 14a). Furthermore, when the RFP^+^ cell density was higher in Injured-GC mice, some ASCs that were traced contacted different RFP^+^ cells continuously and simultaneously before division (Fig. 5f and i1, Supplementary Video 14b). Similar to the Injured-Ctrl group, the maximum consecutive frames that individual RFP^+^ cells contacted ASCs were significantly lower than the total frames that ASCs were traced (Fig. 5i1), indicating that few RFP^+^ cells established constant contact with ASCs before ASC division.

Besides, we could observe more RFP^-^NADH^+^ signal in the injured sites at 1.5 dpi and 2 dpi in the Injured-KO and Injured-GC group before time-lapse imaging than that in the Injured-Ctrl group (Fig. 5a, d-f). These RFP^-^NADH^+^ cells probably included neutrophils, resident macrophages, and fibro/adipogenic progenitors (FAPs), which could regulate MuSCs functionally, but the density of FAPs increased much more slowly than infiltrating neutrophils which peaked around 24 hpi^7,8,10,28,35,76,77^. Therefore, we reasoned that most RFP^-^NADH^+^ cells observed were neutrophils; next, we performed immunostaining in the whole-mount TA muscle using fluorophore-conjugated antibodies of neutrophils (Gr-1 and Ly6G) and found that more than 85% of NADH^+^ cells were Gr-1^+^ or Ly6G^+^ in Injured-GC and Injured-GC mice at 1.5 dpi (Supplementary Figs. 6h-i). ASCs contacted RFP^-^NADH^+^ cells transiently (Fig. 5g2, h2 and i2). In some cases, due to the high density of RFP^-^NADH^+^, ASCs contacted multiple non-myogenic cells simultaneously both in the Injured-KO and Injured-GC groups (Fig. 5d-e). Besides, the total duration of the contact between ASCs and RFP^-^NADH^+^ cells was significantly longer than the contact between ASCs and RFP^+^ cells in the Injured-KO group (Supplementary Fig. 6g). Similarly, continuous contacts with a specific cell before ASC cell division were rarely observed in both Injured-KO and Injured-GC groups. Further, the duration of contact between RFP^+^ MOs/MΦs and ASCs in the Injured-GC group was similar to that in the Injured-Ctrl group, and they were both significantly longer than the contact between non-myogenic cells and ASCs in the Injured-KO group (Supplementary Fig. 6e).

Although the overall proliferation rate of ASCs in the Injured-KO and Injured-GC group was similarly and both slightly impaired by the reduction of macrophages (Fig. 5c and Supplementary Fig. 6j), all ASCs in the three groups could successfully complete cell division at 1.5 dpi, indicating that cell division of ASC did not require continuous contact with a specific RFP^+^ MO/MΦ or other RFP- non-myogenic cells. Notably, we also found that the ASCs mainly migrated along the longitudinal axis (y-axis) of the muscle fiber (Supplementary Figs. 6k-l) and the displacement in the y-axis was significantly larger than in the x and z axes (Supplementary Fig. 6l) in the three groups. After depletion of macrophages, the displacement along the z-axis slightly increased (Supplementary Fig. 6l) but did not exceed the average diameter of a muscle fiber (Supplementary Fig. 6m), which was consistent with the previous study^44^. In addition, the instantaneous migration speed of ASCs in the three groups was similar (Supplementary Fig. 6n). In summary, the depletion of macrophages impaired the proliferation rate but had no influence on the migration speed of the ASCs.

### The role of macrophages during the differentiation stage of muscle regeneration

To further investigate how the impaired recruitment of macrophages influences the differentiation of MuSCs, we performed *in vivo* imaging sessions in the TA muscles at 3, 4, and 15 dpi in the Injured-Ctrl, Injured-GC and Injured-KO groups (Fig. 6). In the Injured-Ctrl group, the RFP^+^ MOs/MΦs filled the injured areas at 3 dpi, started to decrease at 4 dpi, and further decreased at 15 dpi (Fig. 6a, b and e). At 3 dpi and 4 dpi, the RFP^+^ cell density was higher in the Injured-GC group than in the Injured-KO group. In addition, the Injured-GC group had a similar RFP^+^ cell density to the Injured-Ctrl group at 4 dpi (Fig. 6b) presumably because monocyte depletion by drug treatment could increase the local proliferation of macrophage subsets^78^. However, the YFP^+^ area density of Injured-GC and Injured-KO groups was lower than the Injured-Ctrl group at 3-4 dpi (Fig. 6c). In addition, some newly formed myotubes were bifurcated or shrunk with RFP^+^ MOs/MΦs located in their growth path in the Injured-GC and Injured-KO groups at 4 dpi (Fig. 6a). These RFP^+^ MOs/MΦs, which had strong autofluorescence in the YFP channel, may be responsible for the clearance of necrotic muscle fibers^72,73^. Therefore, bifurcation of myotubes may be caused by muscle debris that was not phagocytized in time. At 15 dpi, the injured areas were filled with regenerated myotubes parallel to the intact muscle fibers in all three groups (Fig. 6e). Based on SHG signals, we could also visualize some regenerated blood vessels where an RFP+ monocyte went through (indicated by the arrow in Fig. 6e). Additionally, the SHG signal generated from the sarcomeres of myotubes was stronger and had more apparent periods in the Injured-Ctrl and Injured-GC group, suggesting that myotubes were less mature in the Injured-KO groups (Fig. 6e-f). The SHG signal of some sarcomeres was weaker in the middle of muscle fibers because the nuclei were centrally aligned in the muscle cells (Supplementary Fig. 7a). As expected, the diameter of the myotubes was smaller in the Injured-GC and Injured-KO groups (Fig. 6g). However, there was no significant difference in the density of individual RFP^+^ and YFP^+^ cells between the three groups (Fig. 6h-i). Further, we could observe that some myotubes were bifurcated by intramuscular cells or collagen in the Injured-KO and Injured-GC groups, which never occurred in the Injured-Ctrl group at 15 dpi (Supplementary Fig. 7b). Also, although the collagen density of Injured-KO groups was modestly lower than the Injured-Ctrl groups at 4 dpi, it remained at a high level at 15 dpi (Fig. 6d and Supplementary Fig. 7c), which might be caused by continued fibro/adipogenic progenitor (FAP) accumulation in muscle from Injured-KO groups^79^. Although the RFP^+^ cell density was higher at 4 dpi, the collagen density in the Injured-GC group was similar to that in the Injured-KO group at 15 dpi (Fig. 6a, b, d, Supplementary Fig. 7c), which indicated that the delayed infiltration of RFP^+^ MOs/MΦs at 4 dpi could not reduce the fibrosis at 15 dpi. Together, these data indicated that the macrophage deficiency severely delayed the differentiation of MuSCs and induced intramuscular fibrosis.

**Fig. 6:**
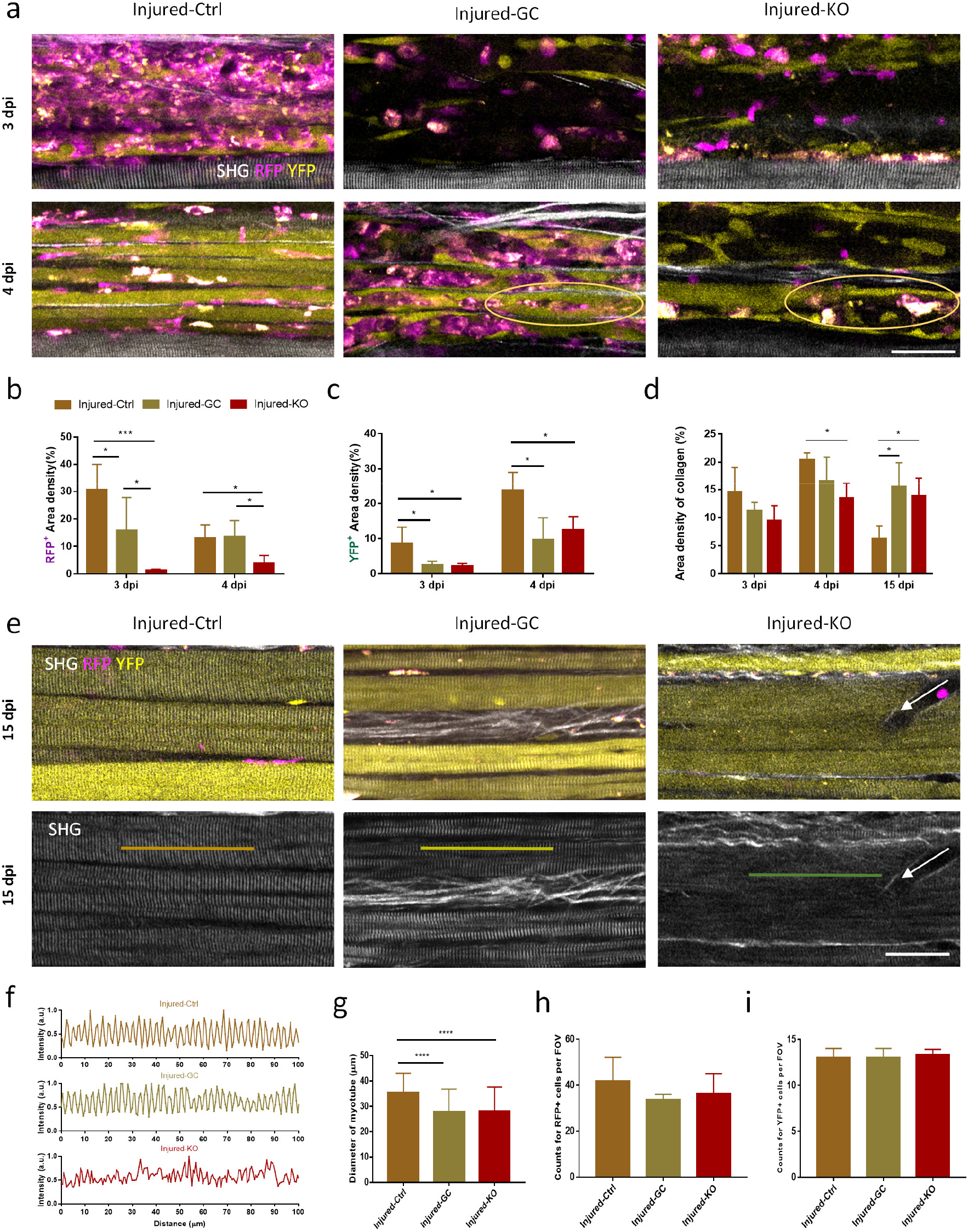
The depletion of macrophages impaired the proliferation and differentiation of MuSCs. (a) Images for the RFP^+^ MOs/MΦs distribution (magenta), YFP^+^ ASCs/myotubes (green) and SHG generated from collagen and muscle fiber (grey) at 3 dpi and 4 dpi in the Injured-Ctrl, Injured-GC, and Injured-KO groups. The bifurcated or shrunk myotubes at 4 dpi in the Injured-GC and Injured-KO groups were delineated by brown circles. (b-d) The statistics of RFP^+^ area (RFP^+^ MOs/MΦs) density (b), YFP^+^ area density (c), area density of collagen (d) at indicated time points after injury of three groups. (e) Representative TPEF and SHG images of newly-regenerated myotubes and RFP^+^ MOs/MΦs at 15 dpi of three groups. Green, YFP^+^ myogenic cells/myotubes; magenta, RFP^+^ MOs/MΦs; grey, SHG generated by muscle fiber and collagen. The arrow indicated a newly-formed blood vessel. (f) The normalized signal profiles along the lines in the e. (g-i) The diameter of newly-regenerated myotubes (g), RFP^+^ cell density (h), and YFP^+^ cell density (i) at 15 dpi of three groups. Error bars were SD. (*: p<0.05; ***: p<0.001; ****: p<0.0001; student t-test). Sale bars: 50 um in (a, e). A.u., arbitrary units.

## Discussion

Using a home-built dual-laser non-linear optical microscope platform, we simultaneously visualized various cell types and both intracellular and extracellular structures in the skeletal muscle, including RFP^+^ monocytes/macrophages, RFP^-^ non-myogenic cells, YFP^+^ MuSCs/ASCs, collagen, and muscle fibers, which facilitates the examination of cellular dynamics of MuSCs and spatiotemporal interaction between non-myogenic and myogenic cells during muscle regeneration. Mitochondrial biogenesis of MuSCs was first observed during activation *in vivo* based on the autofluorescence of the metabolic coenzyme NADH (Fig. 1d). Activated MuSCs are larger in size with more cytoplasm. In previous *in vitro* and *ex vivo* studies, the diameter, the mean forward scatter (FSC) values and the volume of the isolated/stained MuSCs were measured to explore their status^53,54,80,81^. In this study, we first monitored the change of MuSC volume *in vivo* using our two-photon microscope. Consequently, we found that the mitochondrial mass and volume of MuSCs increased synchronously before 2 dpi after injury and returned to basal level at 1 mpi (Fig. 1c and d). This observation was consistent with the previous *in vitro* finding that mitochondrial biogenesis increased continuously in control MuSCs up to 40 hours in culture^45^.

During regeneration, it is believed that non-myogenic cells, primarily infiltrating neutrophils and monocytes/macrophages, interact with MuSCs and regulate their regenerative potential. Previously, it was shown that muscle regeneration was impaired by the genetic ablation of macrophages in the early stages of injury^14,19,20,25^. The proper transition from pro-inflammatory to anti-inflammatory macrophages seems to be required for efficient muscle regeneration^25^. Early studies of *in vitro* co-culture systems also showed that macrophages can protect MuSCs from apoptosis and promote myogenic differentiation^37,82^. In one study, macrophages were found to promote MuSCs activation by secreting ADAMTS1^32^. It was shown that overexpression of ADAMTS1 could increase the ratio of Pax7^+^MyoD^+^ MuSCs in a non-injury state, and macrophages were the primary source of secreted ADAMTS1. However, a detailed examination of how macrophage-MuSC interaction affects MuSCs dynamics and muscle regeneration *in vivo* is still lacking. To achieve this, along with our microscope platform, we introduced two macrophage-depletion models to examine the behavior of macrophages during muscle regeneration. One was Glucocorticoid-treatment (Injured-GC) to suppress the immune response, and the other was the CCR2ko transgenic mouse model (Injured-KO). The data of snap-shot imaging sessions at different time points indicated that the depletion of macrophages did not impair the initial increase or further growth of MuSC volume *in vivo* (Supplementary Fig. 4k). In addition, time-lapse imaging suggested that individual MuSCs without contacting RFP^+^ macrophages in the Injured-KO and Injured-GC group increased in volume to the same extent (~20%) as MuSCs in contact with macrophages in the Injured-Ctrl group. These findings demonstrated that the infiltration of macrophages and direct contact with RFP^+^ monocytes/macrophages were unnecessary for MuSCs activation. In addition, depletion of the neutrophils in the CCR2ko mice did not delay the initial increase in cell volume of MuSCs (Fig. 3d). In summary, the activation of MuSCs may rely more on intrinsic changes in signaling (e.g., loss of Notch signaling) than direct interaction with myeloid cells^40,81,83^.

In contrast to the insignificant role of macrophages in MuSC activation, we found that proliferation and differentiation of MuSCs was impeded by the reduction of macrophages (Fig. 5c, 6c, 6g and Supplementary Fig. 6j), and fibrosis was more significant at 15 dpi in the Injured-KO and Injured-GC groups than that in the Injured-Ctrl group (Fig. 6d and Supplementary Fig. 7c). Therefore, the newly-recruited macrophages promote the proliferation and differentiation of MuSCs but are unnecessary for their activation *in vivo*. However, since MuSC differentiation program requires cell cycle exit after limited rounds of cell division, the balance between MuSC proliferation and subsequent differentiation dictates the regeneration outcome^4^. It is difficult to discern whether macrophage depletion acts on MuSC differentiation directly or indirectly (e.g., through regulating MuSC proliferation) *in vivo*. For *in vitro* coculture studies, contradictory findings regarding macrophages’ role in MuSC differentiation were reported^82,84,85^. This is not surprising, since co-culture setup, cell source and cell state greatly influence the outcome of such experiments. For example, early co-culture studies predominantly utilized macrophage cell line or macrophages stimulated in culture dish to obtain different polarization states rather than macrophages isolated from injured muscle^82,86^. Nevertheless, our findings were in line with previous reports which showed an overall reduction of macrophages impedes satellite cell differentiation during muscle regeneration *in vivo*^14,19,23,25,87^. Combining the autofluorescence of NADH from RFP^-^ non-myogenic cells including resident macrophages, neutrophils and FAPs and TPEF of YFP and RFP, we first demonstrated the mobility of non-myogenic cells and myogenic cells and their spatiotemporal interaction after an acute injury in the mouse model *in vivo* in real-time. Immediately after injury, MuSCs were motionless, but the non-myogenic cells were highly motile. The majority of non-myogenic cells was neutrophils and monocytes/macrophages. Direct cell-cell contacts between MuSCs and non-myogenic cells were mostly frequent and short-lived. Notably, the total number of non-myogenic cells that contacted MuSCs would be overestimated because it could not be verified whether the cells came back after leaving the tracing FOV. Occasionally, we could observe continuous contact between MuSCs and immobile RFP^-^ non-myogenic cells which had a similar spindle shape to MuSCs in the NADH channel (Fig. 3e and Supplementary Fig. 5h-j). These may be the mesenchymal stem cells such as fibro/adipogenic progenitors (FAPs) located quiescently between muscle fibers in the non-injury state and sharing a similar temporal pattern of activation and proliferation with MuSCs after injury^7,76^.

During the proliferation stage, MuSCs/MPs migrated parallel to the intact muscle fiber, slowing down when they entered the M phase and acquired a round shape (Supplementary Video 11-14). At the same time, most non-myogenic cells were also highly dynamic, with their mobility being asynchronous. Therefore, MuSCs could not establish continuous contact with a specific non-myogenic cell before division. This observation differed from previous *in vivo* finding in larval zebrafish^41^. This study observed that MuSCs established pro-longed direct contact with a specific dwelling macrophage before division. In zebrafish, both MuSCs and macrophages were motionless during the proliferation stage of MuSCs, which was conducive to establishing constant contact. However, their communication medium was a pairing between secreted cytokine NAMPT and the chemokine receptor CCR5 but not direct cell-cell signaling. Therefore, non-myogenic cells may mainly regulate the proliferation of MuSCs via paracrine cytokines or metabolite signaling^14,30,31,32,33,34^. As a result, continuous MuSC-non myogenic cell contact is not required for MuSC cell division, albeit the reduction of non-myogenic cell presence in injury sites slows down the overall MuSC proliferation. Intriguingly, the depletion of macrophages which reduced the duration and frequency of contacts between macrophages and myoblasts did not affect instantaneous migration speed of myoblasts and the myoblast still migrated along the longitudinal axis of the muscle fiber (Supplementary Fig. 6k-l), which contrasted the *in vitro* findings that macrophages can promote the migration of MuSCs^82,88,89,90^.

A few limitations in this study should be noted. First, skeletal muscle resident macrophages play an important role during muscle regeneration^43,77^, but we cannot investigate their exact role because the depletion models have no significant impact on the number of resident macrophages^42,65^, and most of them cannot be identified without labeling. Second, NADH signals can, in principle, mark all cell types, but not definitively identify them. Our data showed the interaction between non-myogenic cells and YFP^+^ myogenic cells, but it was still unclear whether different cell types had distinct interaction patterns with myogenic cells. Therefore, transgenic models that have specific markers for different cells are required in further study. Third, we did not study the functional regeneration of muscle fibers after an acute injury in the control and macrophage-depleted groups. The contraction of muscle fibers could be explored by high-speed imaging of second harmony generation (SHG)^91^ and the functional repair of the neuromuscular junctions (a synaptic connection between terminal end of a motor nerve and a muscle)^92,93^ could also be studied further.

In conclusion, we demonstrated a promising method to study the dynamics of different cell types and their interactions in live animals, facilitating further understanding of the mechanism of muscle regeneration on the cellular and tissue level. With this tool in hand, for the first time, we documented MuSC dynamics as well as its interacting pattern with myeloid cells during the course of muscle regeneration in mice. In particular, we found that MuSC-myeloid cell interactions were always transient and prolonged contact was not present nor required for MuSC to undergo quiescence exit or cell division, in contrast with a previously published live imaging study. However, depletion of macrophages impairs muscle regeneration due to moderately reduced MuSC proliferation rate and differentiation. Therefore, we would like to propose a stage-dependent role for myeloid cells to regulate MuSC functions during muscle regeneration. Further studies combining our methods, appropriate transgenic models, or chronic imaging windows^94^ can reveal more about the mechanisms of muscle regeneration and promote the development of therapeutic strategies for muscle diseases such as Duchenne muscular dystrophy (DMD).

## Materials and Methods

### Animal preparation

Pax7^CreERT2(Gaka)^ (Stock No: 017763), Ccr2^RFP/RFP^ (Stock No: 017586) and Rosa-LSL-YFP (Stock No: 006148) mice were from Jackson Laboratory (Bar Harbor, ME, USA). Appropriate mating schemes were used to generate mice to study MuSCs dynamics and interaction between RFP^+^ cells and MuSCs. To enable YFP expression in adult MuSCs, female and male mice, 2-5 months of age, were injected intraperitoneally with 3 doses of tamoxifen (Santa Cruz biotechnology; 200 μg/g of body weight, diluted in corn oil) for 7 days. All experiments were conducted at least 1 week after the last injection. Mice were housed two to four animals per cage with a standard 12-h light/dark cycle in a temperature-controlled environment (22–25 °C with 40–60% humidity) and had *ad libitum* access to food and water. Before the injury, mice were anesthetized by intraperitoneal (i.p.) injection of a ketamine-xylazine mixture (87.5 mg kg^−1^ and 12.5 mg kg^−1^). First, the skin covering the TA muscle was shaved of all fur using depilatory cream (Veet®) and disinfected with 75% EtOH. Next, the TA muscle was punctured at 5 sites with a 25G needle (NN-2522R, Terumo) without incising the skin, and the skin was disinfected again. For cardiotoxin (CTX)-induced injury, 30 ml of 10 mM CTX was injected into TA muscles of anesthetized mice. All the experiments were performed following the guidelines of the Laboratory Animal Facility of the Hong Kong University of Science and Technology (HKUST) and the animal protocols were approved by the Animal Ethics Committee at HKUST.

### Two-photon microscope

The two-photon system is shown in Supplementary Fig. 1, and was modified from the previously reported multimodal nonlinear optical microscope system^72^. In brief, two femtosecond Ti: Sapphire lasers (Chameleon Ultra II, Coherent) with 80MHz were used. One was tuned at 920 nm to excite TPEF of YFP in muscle satellite cells and SHG signals of myosin filament and collagen. The other was tuned at 740 nm to excite TPEF of RFP in macrophages and NADH signals. The power intensity and polarization of the lasers were adjusted by a combination of a half-wave plate (AHWP10M-980, Thorlabs) and a polarizing beamsplitter cube (CCM1-PBS252/M; Thorlabs). The lasers were collimated and expanded by a pair of achromatic doublets to match the 5mm Galvo XY-scan mirror (6215H, Cambridge Technology). Another polarizing beamsplitter cube was chosen to combine the two lasers and then direct them to the Galvo X scanner, which enables any combination of the two different wavelengths of laser (920 & 740 nm, 810 & 740 nm, or 920 & 810 nm). The XY scanners were mutually conjugated through a *4f* relay formed by L5 and L6, both of which consist of two doublets (49-392, Edmunds). The Galvo Y and the rear pupil of the water-immersive objective (OL: XLPLN25XSVMP2, 25×, 1.05 NA, Olympus) were then conjugated by the scan lens L7 (two doublets, 49-392, Edmunds) and the tube lens L8 (49-365, Edmund) operating in the *4f* relay configuration. The objective was mounted on a motorized linear actuator (LNR50SEK1, Thorlabs) for axial sectioning. The epifluorescence collected by the objective was reflected by a 705 nm long-pass dichroic mirror (D1: FF705-Di01-25×36, Semrock) and directed to the detection unit. In addition, a half-wave plate (AHWP10M-980, Thorlabs) was applied before the D1 to change the polarization of the exciting laser and selectively strengthen the SHG signal of collagen or sarcomere. The fluorescence was further separated by a second dichroic mirror (D2: FF495-Di03-25×36, Semrock) and directed into two current photomultiplier modules (PMT: H11461-03 and H11461-01, Hamamatsu). Two band-pass filters F1 (FF02-447/60 or FF01-357/44, Semrock) and F2 (FF03-525/50 or FF01-593/46, Semrock) and two short-pass filters (FF01-680SP, Semrock) were placed before PMTs respectively to reject the excitation laser and select the fluorescence. The output currents of both PMTs were then converted to the voltage by two current amplifiers (SR570, Stanford Research and DLPCA-200, Femto) and subsequently fed into a multifunction data acquisition device (PCIe-6353, National Instrument). Custom-written C# software running in Visual Studio (Microsoft) was used to control all the hardware and acquire the TPEF images.

### *In vivo* imaging

The mice were anesthetized with a ketamine/xylazine mixture before surgery. The skin covering the TA muscle was resected to expose approximately 0.5 cm^2^ of the muscle. The fascia and epimysium were gently removed to avoid damaging the muscle fibers. Next, the hind limb was put in the middle of a customized stage consisting of metal and heat-insulated materials, as shown in Supplementary Fig. 3. Both sides of the stage were wrapped in a soft heating plate. The coverslip (Ø=18 mm) attached to a metal support holder was put over the muscle lightly but firmly to minimize the pressure applied to the muscles. The stage securing the mouse was placed on a five-axis stage beneath our prototype microscope. The five-axis stage allows three-axis translation and ±5° pitch and roll flexure motion. The surface of the coverslip was aligned perpendicular to the objective axis by adjusting the roll and pitch angles of the stage guided by the reflection of a laser diode (CPS635R, Thorlabs) on the coverslip. A laser tuned to 920nm and another to 740nm were switched sequentially to obtain the YFP/SHG and RFP/NADH signals, respectively. For snap-shot imaging, the excitation laser power at the sample plane was less than 35mW, and the image integration time was less than 4s.

For several hours of time-lapse imaging, the laser power at 740nm was less than 25 mW, and the image integration time for a single frame was less than 2s to avoid photo-bleaching and photodamage. Serial optical sections were captured at 3 um steps to a total depth of 60-90 um every 5 mins. The scanning field of view (FOV) of the injured site was 300 μm × 300 μm/400 μm × 400 μm (256*256 pixels or 512*512 pixels). Besides, the laser power of 740nm was higher in the time-lapse imaging of ASCs at the proliferation stage. To measure the volume of single muscle satellite cells, the YFP signal of MuSCs (100 μm× 100 μm, 256*256 pixels or 50 μm× 50 μm, 128*128 pixels) was captured with 1 μm steps to cover entire cells. During time-lapse imaging, the mice were anesthetized with 0.5-1% isoflurane. A heating pad beneath the mice was set to 36°C to maintain their body temperature. In addition, the muscle temperature was maintained at 28-30°C using a heating plate wrapping the stage holding the muscle and the lens warmer enveloping the objective.

### *In situ* staining of cells for *in vivo* imaging

Mitotracker deep red (MTDR: MitoTracker™ Deep Red FM, ThermoFisher) was dissolved in anhydrous dimethylsulfoxide (DMSO) to a final concentration of 1mM and then diluted to 500nM using saline or Phosphate-buffered saline (PBS). The dye was applied topically to the exposed TA muscle. Approximately 30 minutes after staining, saline or PBS was applied to the muscle to remove the unwanted dye, and the *in vivo* imaging started. One laser was tuned to 810 nm, and the F2 was switched to another band-pass filter (FF01_650/60-25, Semrock) to obtain the TPEF image of MDTR.

### Neutrophil depletion

Pax7^CreER/+^/Rosa-YFP/CCR2^RFP/+^ mice were intraperitoneally treated with 250 μg anti-mouse ly6G antibody (clone 1A8, Bio X Cell) diluted in 150 μL saline 1 day before needle injury to deplete neutrophils *in vivo*. In addition, the Pax7^CreER/+^/Rosa-YFP/CCR2^RFP/RFP^ mice were intraperitoneally injected with cyclophosphamide (CP: 6055-19-2, MedChemExpress) in two doses to induce neutropenia in the CCR2-deficient mice^70^. Initially, 150 mg kg-1 was administered in 200 μl saline as the first dose on day 1, and the second dose of 100 mg kg–1 was administered on day 4. The mice were injured 12 hours after the last injection.

### Glucocorticoid treatment to deplete macrophages

Glucocorticoid treatment was conducted as previously described^24^. Briefly, Pax7^CreER^/Rosa-YFP/CCR2^RFP/+^ mice were treated with 0.9% saline drinking water supplemented with 100 μg/ml corticosterone (Sigma, St.Louis, MO) starting from 3 days before needle injury until the mice were imaged at specific time points after the injury. The water was changed daily, and water consumption was monitored. There was no difference between the treated mice and the control mice (around 7-10 ml/day) during the experiments.

### Whole-mount muscle immunostaining

The tibialis anterior muscles were dissected at specific time points after needle injury, fixed in 4% PFA for 3-4 h at 4 °C on a shaker and rinsed 3 times with PBS. 0.1 M glycine PBS solution was used for blocking free aldehydes. Tissue was then permeabilized using 0.5% Triton X in PBS for 3-4 h and blocked with blocking buffer (0.5% Triton X-100 in PBS containing 4% BSA) for 3-4 h. Samples were incubated with primary antibodies and subsequently, with secondary antibodies diluted in blocking buffer overnight at 4 °C. Washing was performed between steps by rinsing the samples 3 times with 0.3% Triton X in PBS for 1 h on a shaker. The samples were placed in Mowiol mounting medium and examined with two-photon fluorescence microscopy. The primary antibodies used are from SouthernBiotech (anti-Type I collagen) and Biolegend (anti-CD45, anti-Ly6G, anti-Gr-1 and anti-CD11b). Secondary antibodies are from the Jackson ImmunoResearch (Alexa-488/594/647 anti-rat IgG and Alexa-488 anti-goat IgG).

### Imaging processing and analysis

The images were processed with MATLAB or ImageJ^95^.

#### Cell volume and mitochondria mass measurement of MuSCs

After smoothing and thresholding the 3D-sectioning stacks, the volume was calculated using the plugin “3D Objects Counter” of ImageJ. The threshold was set to the fluorescent intensity covering the whole cell in the stacks’ maximum-intensity projection (MIP) image. The NADH signal of a MuSC was captured at the best focal plane of the cell. The YFP signal at that plane was regarded as a mask to depict the cell boundary. The mitochondrial mass was calculated as the area size of the NADH signal inside the mask after smoothing, thresholding and binarizing using ImageJ ‘Threshold’ plugin.

#### Cell density calculation

For myogenic cell density, YFP^+^ were counted for the MIP of every FOV (300 um*300 um*60 um) before and at 2 dpi. For macrophage density, RFP^+^ cells were counted for every FOV before and at 12 hpi. When YFP^+^ cells started to differentiate or RFP^+^ cell density was so high that individual cells could not be identified, we calculated the YFP^+^/RFP^+^ area density (%) in the injured sites as the YFP^+^/RFP^+^ cell density. The density was calculated as the ratio of the YFP^+^ or RFP^+^ area to the injured site indicated by SHG (excluding the intact muscle fiber areas based on periodic signals). However, some RFP^+^ macrophages had strong autofluorescence in the YFP channel after 1 dpi, which would interfere with the accuracy of YFP^+^ area density. Therefore, we merged the YFP stacks with the RFP stacks to create RGB stacks. Then we selected the green (YFP^+^RFP^-^) signal using a color threshold and calculated the YFP^+^ area density in Matlab. The cell density for each mouse was the average value of 3-6 FOVs.

#### Collagen density calculation

To calculate the collagen density (%), we projected the stacks of SHG channels every three slices for 15 slices and manually depicted the dark zone of the injured site where there was no SHG signal from muscle fibers. The contrast of the SHG of collagen in the dark zone was adjusted, and the images were median filtered (Neighborhood size was 6*3). The density was calculated as the ratio of the collagen area to the dark zone area after thresholding using multithresh.m in Matlab. The collagen density of one FOV was the average value of the slices, and the collagen density for one mouse was the average value of 3-6 FOVs.

#### Time-lapse imaging processing

Stacks of 4 channels from optical sections were merged, converted to hyperstacks and aligned by refrence to the SHG signal using the plugin ‘Correct 3D drift’ in ImageJ. To calculate the minimal distance of individual MuSCs from macrophages from 4.5 to 12 hpi, RFP^+^ and YFP^+^ stacks (volume is 300 um*300 um*60 um) for different frames were first denoised, set the gama value as 0.8 and smoothed, then converted to binary images using ImageJ ‘Threshold’ plugin with the ‘Default’ auto thresholding method. The 3D distance matrix was calculated from the binary image of macrophages using bwdistsc.m^96^ in the Matlab so that the further a point was from macrophages, the higher the pixel value. The area of each YFP^+^ cell was labeled using bwlabeln.m in Matlab. The minimal distance of every YFP^+^ cell from macrophages was the minimal value of the corresponding pixels in the distance matrix. If the minimum distance was 0 μm, the YFP^+^ cells were in contact with macrophages. In addition, to analyze the contact between YFP^+^ cells and RFP^+^ cells at the proliferation stage, the slices covering the whole YFP^+^ cell were stacked as a maximum z projection. The images were then denoised and smoothed. We tracked individual RFP^+^ that contacted YFP^+^ ASCs and ASCs using the ImageJ plugin ‘Manual Tracking.’ Direct cell-cell contact was defined by at least one RFP^+^ positive pixel juxtaposed to at least one YFP^+^ pixels and was confirmed by the examination of individual z planes in the raw stacks. The analysis of the contact between YFP^+^ cells and RFP^-^NADH^+^ non-myogenic cells was similar. Because the difference between the cell boundaries depicted by the NADH signal and RFP/YFP signal was less than 2.5 μm laterally and 4 μm axially, we projected two more slices in two directions for YFP^+^ cells, and if the lateral distance between YFP^+^ cells and RFP^-^NADH^+^ non-myogenic cells was less than 2.5 μm, contact was established. Notably, only the cells with visible NADH signal (mainly located within 20-30 um in depth) were analyzed. In addition, the image sequences after registration were further analyzed in the Imaris (Bitplane AG, Zurich, Switzerland) to obtain the migration statistics (the migration speed, migration direction and so on).

### Statistical analysis

Statistical analysis and data visualization were performed using GraphPad Prism 7 software. All similar measurements were performed on more than three mice. Data are presented as mean ± s.d. and α = 0.05 for all analyses.

## References

1. Shadrin, I. Y., Khodabukus, A. & Bursac, N. Striated muscle function, regeneration, and repair. Cellular and molecular life sciences 73, 4175–4202 (2016).

2. Mauro, A. Satellite cell of skeletal muscle fibers. The Journal of biophysical and biochemical cytology 9, 493 (1961).

3. Wang, Y. X. & Rudnicki, M. A. Satellite cells, the engines of muscle repair. Nature reviews Molecular cell biology 13, 127–133 (2012).

4. Yin, H., Price, F. & Rudnicki, M. A. Satellite cells and the muscle stem cell niche. Physiol. Rev. 93, 23–67 (2013).

5. Chen, X. et al. Lockd promotes myoblast proliferation and muscle regeneration via binding with DHX36 to facilitate 5’ UTR rG4 unwinding and Anp32e translation. Cell Reports 39, 110927 (2022).

6. Sambasivan, R. et al. Pax7-expressing satellite cells are indispensable for adult skeletal muscle regeneration. Development 138, 3647–3656 (2011).

7. Murphy, M. M., Lawson, J. A., Mathew, S. J., Hutcheson, D. A. & Kardon, G. Satellite cells, connective tissue fibroblasts and their interactions are crucial for muscle regeneration. Development 138, 3625–3637 (2011).

8. Wosczyna, M. N. & Rando, T. A. A muscle stem cell support group: coordinated cellular responses in muscle regeneration. Developmental cell 46, 135–143 (2018).

9. De Micheli, A. J. et al. Single-cell analysis of the muscle stem cell hierarchy identifies heterotypic communication signals involved in skeletal muscle regeneration. Cell reports 30, 3583–3595. e5 (2020).

10. Oprescu, S. N., Yue, F., Qiu, J., Brito, L. F. & Kuang, S. Temporal dynamics and heterogeneity of cell populations during skeletal muscle regeneration. IScience 23, 100993 (2020).

11. Tidball, J. G. Regulation of muscle growth and regeneration by the immune system. Nature Reviews Immunology 17, 165–178 (2017).

12. Chazaud, B. Inflammation during skeletal muscle regeneration and tissue remodeling: application to exercise-induced muscle damage management. Immunol. Cell Biol. 94, 140–145 (2016).

13. Gordon, S. & Taylor, P. R. Monocyte and macrophage heterogeneity. Nature reviews immunology 5, 953–964 (2005).

14. Arnold, L. et al. Inflammatory monocytes recruited after skeletal muscle injury switch into antiinflammatory macrophages to support myogenesis. J. Exp. Med. 204, 1057–1069 (2007).

15. Varga, T. et al. Highly dynamic transcriptional signature of distinct macrophage subsets during sterile inflammation, resolution, and tissue repair. The Journal of Immunology 196, 4771–4782 (2016).

16. Tacke, F. et al. Immature monocytes acquire antigens from other cells in the bone marrow and present them to T cells after maturing in the periphery. J. Exp. Med. 203, 583–597 (2006).

17. Dal-Secco, D. et al. A dynamic spectrum of monocytes arising from the in situ reprogramming of CCR2 monocytes at a site of sterile injury. J. Exp. Med. 212, 447–456 (2015).

18. Novak, M. L., Weinheimer-Haus, E. M. & Koh, T. J. Macrophage activation and skeletal muscle healing following traumatic injury. J. Pathol. 232, 344–355 (2014).

19. Martinez, C. O. et al. Regulation of skeletal muscle regeneration by CCR2-activating chemokines is directly related to macrophage recruitment. American Journal of Physiology-Regulatory, Integrative and Comparative Physiology 299, R832–R842 (2010).

20. Lu, H. et al. Macrophages recruited via CCR2 produce insulin-like growth factor-1 to repair acute skeletal muscle injury. The FASEB Journal 25, 358–369 (2011).

21. Arnold, L. et al. CX3CR1 deficiency promotes muscle repair and regeneration by enhancing macrophage ApoE production. Nature communications 6, 1–12 (2015).

22. Summan, M. et al. Macrophages and skeletal muscle regeneration: a clodronate-containing liposome depletion study. American Journal of Physiology-Regulatory, Integrative and Comparative Physiology 290, R1488–R1495 (2006).

23. Segawa, M. et al. Suppression of macrophage functions impairs skeletal muscle regeneration with severe fibrosis. Exp. Cell Res. 314, 3232–3244 (2008).

24. Gao, Y., Li, Y., Guo, X., Wu, Z. & Zhang, W. Loss of STAT1 in bone marrow-derived cells accelerates skeletal muscle regeneration. PLoS One 7, e37656 (2012).

25. Wang, H. et al. Altered macrophage phenotype transition impairs skeletal muscle regeneration. The American journal of pathology 184, 1167–1184 (2014).

26. Teixeira, C. d. F. P. et al. Neutrophils do not contribute to local tissue damage, but play a key role in skeletal muscle regeneration, in mice injected with Bothrops asper snake venom. Muscle & Nerve: Official Journal of the American Association of Electrodiagnostic Medicine 28, 449–459 (2003).

27. Kawanishi, N., Mizokami, T., Niihara, H., Yada, K. & Suzuki, K. Neutrophil depletion attenuates muscle injury after exhaustive exercise. Med. Sci. Sports Exerc. 48, 1917–1924 (2016).

28. Seo, B. R. et al. Skeletal muscle regeneration with robotic actuation–mediated clearance of neutrophils. Science translational medicine 13, eabe8868 (2021).

29. Koike, H., Manabe, I. & Oishi, Y. Mechanisms of cooperative cell-cell interactions in skeletal muscle regeneration. Inflammation and Regeneration 42, 1–11 (2022).

30. Tidball, J. G. & Villalta, S. A. Regulatory interactions between muscle and the immune system during muscle regeneration. American Journal of Physiology-Regulatory, Integrative and Comparative Physiology 298, R1173–R1187 (2010).

31. Furrer, R., Eisele, P. S., Schmidt, A., Beer, M. & Handschin, C. Paracrine cross-talk between skeletal muscle and macrophages in exercise by PGC-1α-controlled BNP. Scientific reports 7, 1–12 (2017).

32. Du, H. et al. Macrophage-released ADAMTS1 promotes muscle stem cell activation. Nature communications 8, 1–11 (2017).

33. Shang, M. et al. Macrophage-derived glutamine boosts satellite cells and muscle regeneration. Nature 587, 626–631 (2020).

34. Girardi, F. et al. TGFβ signaling curbs cell fusion and muscle regeneration. Nature communications 12, 1–16 (2021).

35. Larouche, J. A. et al. Neutrophil and natural killer cell imbalances prevent muscle stem cell–mediated regeneration following murine volumetric muscle loss. Proceedings of the National Academy of Sciences 119, e2111445119 (2022).

36. Sonnet, C. et al. Human macrophages rescue myoblasts and myotubes from apoptosis through a set of adhesion molecular systems. J. Cell. Sci. 119, 2497–2507 (2006).

37. Chazaud, B. et al. Satellite cells attract monocytes and use macrophages as a support to escape apoptosis and enhance muscle growth. J. Cell Biol. 163, 1133–1143 (2003).

38. Ceafalan, L. C. et al. Skeletal muscle regeneration involves macrophage-myoblast bonding. Cell adhesion & migration 12, 228–235 (2018).

39. Choo, H., Canner, J. P., Vest, K. E., Thompson, Z. & Pavlath, G. K. A tale of two niches: differential functions for VCAM-1 in satellite cells under basal and injured conditions. American Journal of Physiology-Cell Physiology (2017).

40. Kann, A. P., Hung, M. & Krauss, R. S. Cell–cell contact and signaling in the muscle stem cell niche. Curr. Opin. Cell Biol. 73, 78–83 (2021).

41. Ratnayake, D. et al. Macrophages provide a transient muscle stem cell niche via NAMPT secretion. Nature 591, 281–287 (2021).

42. Wang, X. et al. Heterogeneous origins and functions of mouse skeletal muscle-resident macrophages. Proceedings of the National Academy of Sciences 117, 20729–20740 (2020).

43. Babaeijandaghi, F. et al. Metabolic reprogramming of skeletal muscle by resident macrophages points to CSF1R inhibitors as muscular dystrophy therapeutics. Science Translational Medicine 14, eabg7504 (2022).

44. Webster, M. T., Manor, U., Lippincott-Schwartz, J. & Fan, C. Intravital imaging reveals ghost fibers as architectural units guiding myogenic progenitors during regeneration. Cell stem cell 18, 243–252 (2016).

45. Zhou, S. et al. Paxbp1 controls a key checkpoint for cell growth and survival during early activation of quiescent muscle satellite cells. Proceedings of the National Academy of Sciences 118, e2021093118 (2021).

46. Li, D., Zheng, W. & Qu, J. Y. Time-resolved spectroscopic imaging reveals the fundamentals of cellular NADH fluorescence. Opt. Lett. 33, 2365–2367 (2008).

47. Quinn, K. P. et al. Quantitative metabolic imaging using endogenous fluorescence to detect stem cell differentiation. Scientific reports 3, 1–10 (2013).

48. Mellem, D. et al. Fragmentation of the mitochondrial network in skin in vivo. PLoS One 12, e0174469 (2017).

49. Plotnikov, S. V., Millard, A. C., Campagnola, P. J. & Mohler, W. A. Characterization of the myosin-based source for second-harmonic generation from muscle sarcomeres. Biophys. J. 90, 693–703 (2006).

50. Sun, Q. et al. Label-free multimodal nonlinear optical microscopy reveals fundamental insights of skeletal muscle development. Biomedical optics express 5, 158–166 (2014).

51. Chen, X., Nadiarynkh, O., Plotnikov, S. & Campagnola, P. J. Second harmonic generation microscopy for quantitative analysis of collagen fibrillar structure. Nature protocols 7, 654–669 (2012).

52. Anderson, J. E. A role for nitric oxide in muscle repair: nitric oxide–mediated activation of muscle satellite cells. Mol. Biol. Cell 11, 1859–1874 (2000).

53. Rodgers, J. T. et al. mTORC1 controls the adaptive transition of quiescent stem cells from G0 to GAlert. Nature 510, 393–396 (2014).

54. Fukada, S. et al. Molecular signature of quiescent satellite cells in adult skeletal muscle. Stem Cells 25, 2448–2459 (2007).

55. Rocheteau, P., Gayraud-Morel, B., Siegl-Cachedenier, I., Blasco, M. A. & Tajbakhsh, S. A subpopulation of adult skeletal muscle stem cells retains all template DNA strands after cell division. Cell 148, 112–125 (2012).

56. Evano, B. & Tajbakhsh, S. Skeletal muscle stem cells in comfort and stress. NPJ Regenerative medicine 3, 1–13 (2018).

57. l’Honoré, A. et al. The role of Pitx2 and Pitx3 in muscle stem cells gives new insights into P38α MAP kinase and redox regulation of muscle regeneration. Elife 7, e32991 (2018).

58. Gailhouste, L. et al. Fibrillar collagen scoring by second harmonic microscopy: a new tool in the assessment of liver fibrosis. J. Hepatol. 52, 398–406 (2010).

59. Liu, X. et al. Type I collagen promotes the migration and myogenic differentiation of C2C12 myoblasts via the release of interleukin-6 mediated by FAK/NF-κB p65 activation. Food & function 11, 328–338 (2020).

60. Tiaho, F., Recher, G. & Rouède, D. Estimation of helical angles of myosin and collagen by second harmonic generation imaging microscopy. Optics express 15, 12286–12295 (2007).

61. Saederup, N. et al. Selective chemokine receptor usage by central nervous system myeloid cells in CCR2-red fluorescent protein knock-in mice. PloS one 5, e13693 (2010).

62. Dearth, C. L. et al. Skeletal muscle cells express ICAM-1 after muscle overload and ICAM-1 contributes to the ensuing hypertrophic response. PLoS One 8, e58486 (2013).

63. Lau, J. et al. Intravital multiphoton imaging of mouse tibialis anterior muscle. Intravital 5, e1156272 (2016).

64. Heinonen, I. et al. Local heating, but not indirect whole body heating, increases human skeletal muscle blood flow. J. Appl. Physiol. 111, 818–824 (2011).

65. Russo-Marie, F. Macrophages and the glucocorticoids. J. Neuroimmunol. 40, 281–286 (1992).

66. Pan, D. et al. The CCL2/CCR2 axis is critical to recruiting macrophages into acellular nerve allograft bridging a nerve gap to promote angiogenesis and regeneration. Exp. Neurol. 331, 113363 (2020).

67. Seo, B. R. et al. Skeletal muscle regeneration with robotic actuation–mediated clearance of neutrophils. Science translational medicine 13, eabe8868 (2021).

68. Bruhn, K. W., Dekitani, K., Nielsen, T. B., Pantapalangkoor, P. & Spellberg, B. Ly6G-mediated depletion of neutrophils is dependent on macrophages. Results in immunology 6, 5–7 (2016).

69. Stackowicz, J., Jönsson, F. & Reber, L. L. Mouse models and tools for the in vivo study of neutrophils. Frontiers in immunology 10, 3130 (2020).

70. Katkar, G. D. et al. NETosis and lack of DNase activity are key factors in Echis carinatus venom-induced tissue destruction. Nature communications 7, 1–13 (2016).

71. Zhang, J. et al. CD8 T cells are involved in skeletal muscle regeneration through facilitating MCP-1 secretion and Gr1high macrophage infiltration. The Journal of Immunology 193, 5149–5160 (2014).

72. Qin, Z. et al. In vivo two-photon imaging of macrophage activities in skeletal muscle regeneration (Imaging, Manipulation, and Analysis of Biomolecules, Cells, and Tissues XVI Ser. 10497, SPIE, 2018).

73. Riew, T. et al. Progressive accumulation of autofluorescent granules in macrophages in rat striatum after systemic 3-nitropropionic acid: a correlative light-and electron-microscopic study. Histochem. Cell Biol. 148, 517–528 (2017).

74. Chang, W., Yang, Y., Lu, H., Li, I. & Liau, I. Spatiotemporal characterization of phagocytic NADPH oxidase and oxidative destruction of intraphagosomal organisms in vivo using autofluorescence imaging and Raman microspectroscopy. J. Am. Chem. Soc. 132, 1744–1745 (2010).

75. Tabula Muris Consortium. Single-cell transcriptomics of 20 mouse organs creates a Tabula Muris. Nature 562, 367–372 (2018).

76. Wosczyna, M. N. et al. Mesenchymal stromal cells are required for regeneration and homeostatic maintenance of skeletal muscle. Cell reports 27, 2029–2035. e5 (2019).

77. Brigitte, M. et al. Muscle resident macrophages control the immune cell reaction in a mouse model of notexin-induced myoinjury. Arthritis & Rheumatism: Official Journal of the American College of Rheumatology 62, 268–279 (2010).

78. Côté, C. H., Bouchard, P., van Rooijen, N., Marsolais, D. & Duchesne, E. Monocyte depletion increases local proliferation of macrophage subsets after skeletal muscle injury. BMC Musculoskeletal Disorders 14, 1–11 (2013).

79. Lemos, D. R. et al. Nilotinib reduces muscle fibrosis in chronic muscle injury by promoting TNF-mediated apoptosis of fibro/adipogenic progenitors. Nat. Med. 21, 786–794 (2015).

80. Wang, G. et al. p110α of PI3K is necessary and sufficient for quiescence exit in adult muscle satellite cells. EMBO J. 37, e98239 (2018).

81. Verma, M. et al. Muscle satellite cell cross-talk with a vascular niche maintains quiescence via VEGF and notch signaling. Cell stem cell 23, 530–543. e9 (2018).

82. Cantini, M. et al. Macrophages regulate proliferation and differentiation of satellite cells. Biochem. Biophys. Res. Commun. 202, 1688–1696 (1994).

83. Goel, A. J., Rieder, M., Arnold, H., Radice, G. L. & Krauss, R. S. Niche cadherins control the quiescence-to-activation transition in muscle stem cells. Cell reports 21, 2236–2250 (2017).

84. Cantini, M. et al. Macrophage-secreted myogenic factors: a promising tool for greatly enhancing the proliferative capacity of myoblasts in vitro and in vivo. Neurological Sciences 23, 189–194 (2002).

85. Merly, F., Lescaudron, L., Rouaud, T., Crossin, F. & Gardahaut, M. F. Macrophages enhance muscle satellite cell proliferation and delay their differentiation. Muscle Nerve 22, 724–732 (1999).

86. Saclier, M. et al. Differentially activated macrophages orchestrate myogenic precursor cell fate during human skeletal muscle regeneration. Stem Cells 31, 384–396 (2013).

87. Tidball, J. G. & Wehling-Henricks, M. Macrophages promote muscle membrane repair and muscle fibre growth and regeneration during modified muscle loading in mice in vivo. J. Physiol. (Lond.) 578, 327–336 (2007).

88. Chazaud, B. et al. Dual and beneficial roles of macrophages during skeletal muscle regeneration. Exerc. Sport Sci. Rev. 37, 18–22 (2009).

89. Lesault, P. et al. Macrophages improve survival, proliferation and migration of engrafted myogenic precursor cells into MDX skeletal muscle. (2012).

90. Venter, C. & Niesler, C. A triple co-culture method to investigate the effect of macrophages and fibroblasts on myoblast proliferation and migration. BioTechniques 64, 52–58 (2018).

91. Mercier, L. et al. In vivo imaging of skeletal muscle in mice highlights muscle defects in a model of myotubular myopathy. Intravital 5, e1168553 (2016).

92. de Paiva, A., Meunier, F. A., Molgó, J., Aoki, K. R. & Dolly, J. O. Functional repair of motor endplates after botulinum neurotoxin type A poisoning: biphasic switch of synaptic activity between nerve sprouts and their parent terminals. Proceedings of the National Academy of Sciences 96, 3200–3205 (1999).

93. Bermedo-García, F., Zelada, D., Martínez, E., Tabares, L. & Henríquez, J. P. Functional regeneration of the murine neuromuscular synapse relies on long-lasting morphological adaptations. BMC biology 20, 1–18 (2022).

94. Jacquemin, G. et al. Longitudinal high-resolution imaging through a flexible intravital imaging window. Science Advances 7, eabg7663 (2021).

95. Schindelin, J. et al. Fiji: an open-source platform for biological-image analysis. Nature methods 9, 676–682 (2012).

96. Mishchenko, Y. 3d euclidean distance transform for variable data aspect ratio. MATLAB Central File Exchange, September (2012).

