## Supplementary Info for "Intravital microscopy of satellite cell dynamics and their interaction with myeloid cells during skeletal muscle regeneration"

### Supplementary Figures

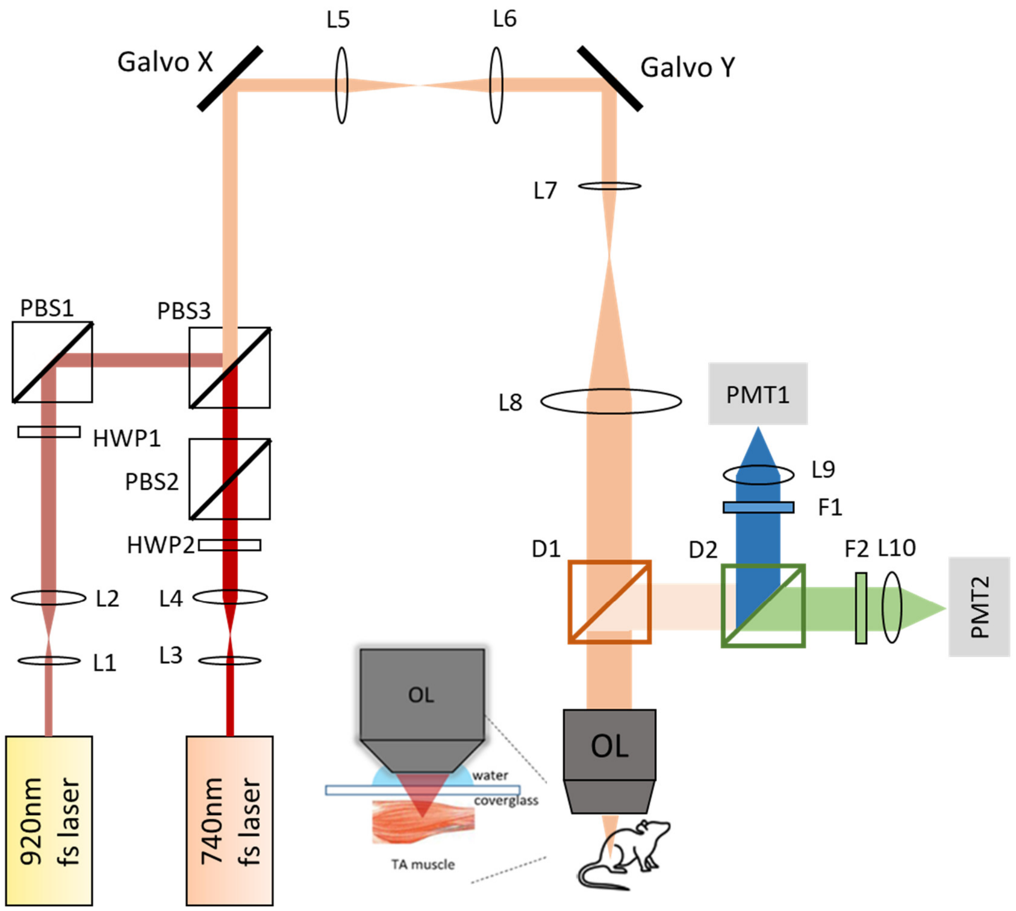


**Supplementary Fig. 1: Schematic diagram of our two-photon microscope system for *in vivo* muscle imaging.** L1-L10: lenses; HWP1-2: half-wave plates; PBS1-3: polarizing beamsplitter cubes; D1-D2: dichroic mirrors; OL: Objective Lens; F1-F2: filters; PMT1-2: photomultiplier tubes.

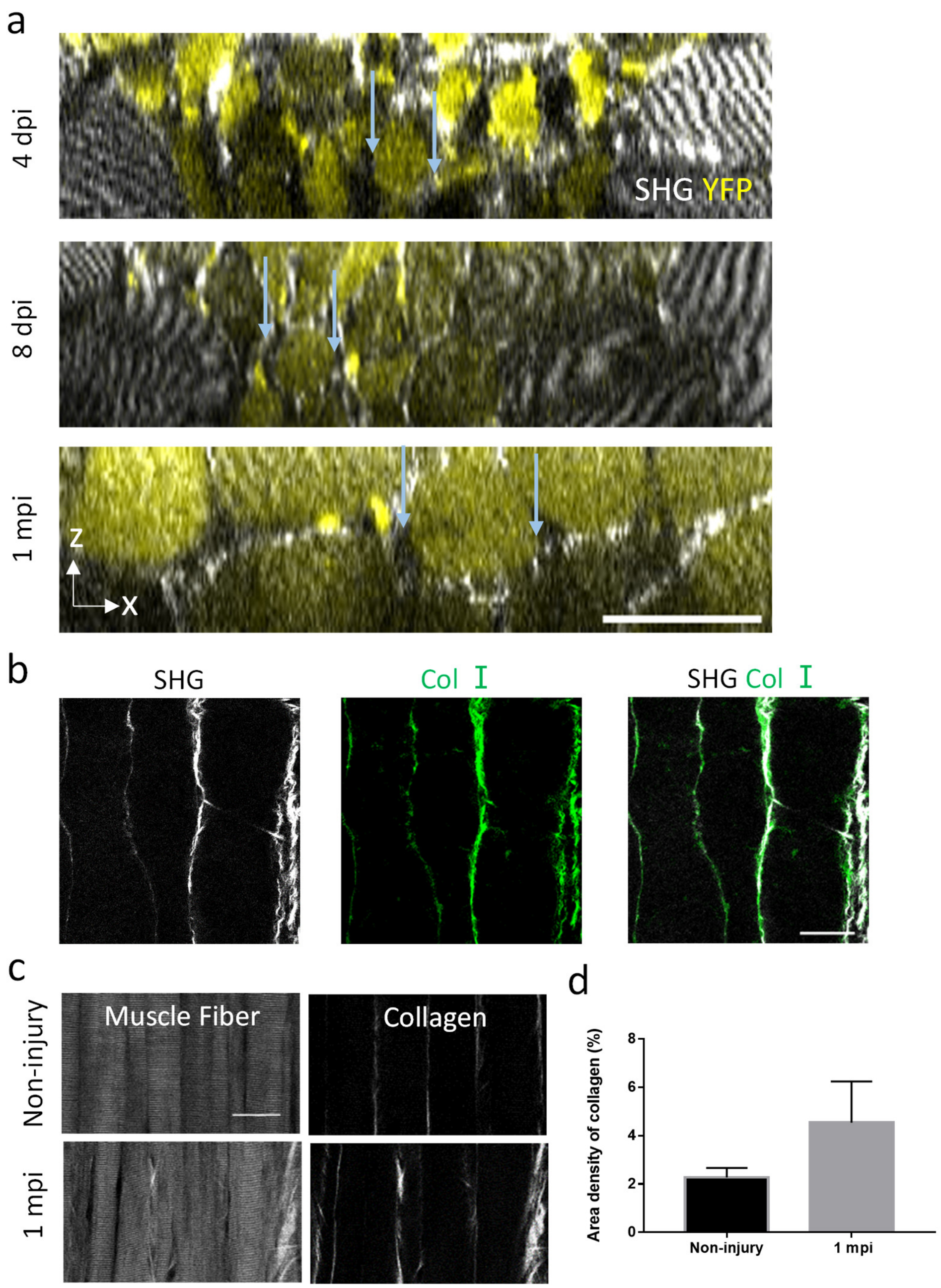


**Supplementary Fig. 2, related to Fig. 1. The SHG signal could reveal the collagen content at the injured sites during muscle regeneration.** (a) The TPEF and SHG images of the x-z cross-section of the injured site at 4 dpi, 8 dpi and 1 mpi. The newly-form myotubes (YFP, yellow) were surrounded by the collagen (SHG, grey), which was indicated by the blue arrows. (b) Whole-mount muscle immunostaining of 1 dpi TA muscle imaged by SHG from collagen (SHG, grey) and fluorescence from Collagen I (Col I, green). (c) Representative images of the SHG from muscle fibers (left) and collagen (right) which were enhanced by adjusting the polarization of exciting laser at non-injury and 1 mpi. (d) The area density of collagen (SHG) at Non-injury and 1 mpi. (n =3 mice per groups). Scale bars: 60 um in a, 50 um in b-c. Error bars, SD, t-test.

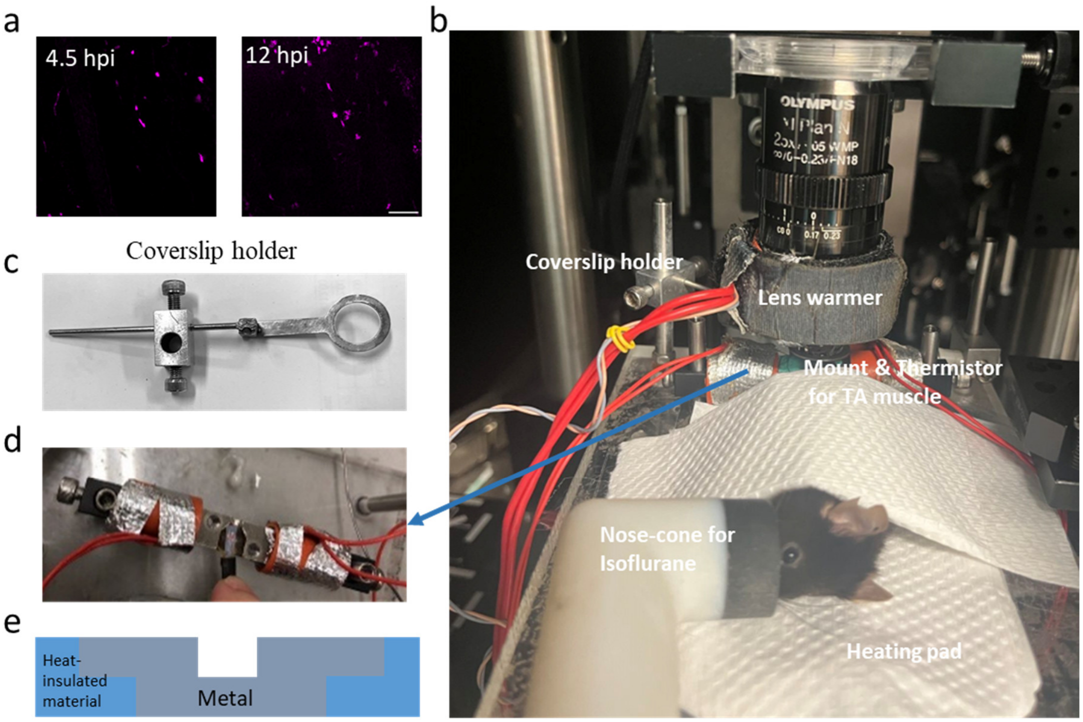


**Supplementary Fig. 3: Mouse positioned for *in vivo* two-photon imaging.** (a) Image sequences of RFP$^{+}$ monocytes/macrophages in the TA muscle without heating for the leg from 4.5 to 12 hpi. Scale bar: 50 um. (b) Stage setup. (c) Coverslip holder. (d-e) Detailed design for the stage securing the tibialis anterior muscle. The stage consisted of three parts, two were made from heat-insulating material and the other of metal. The thermistor wrapped both sides of the stage.

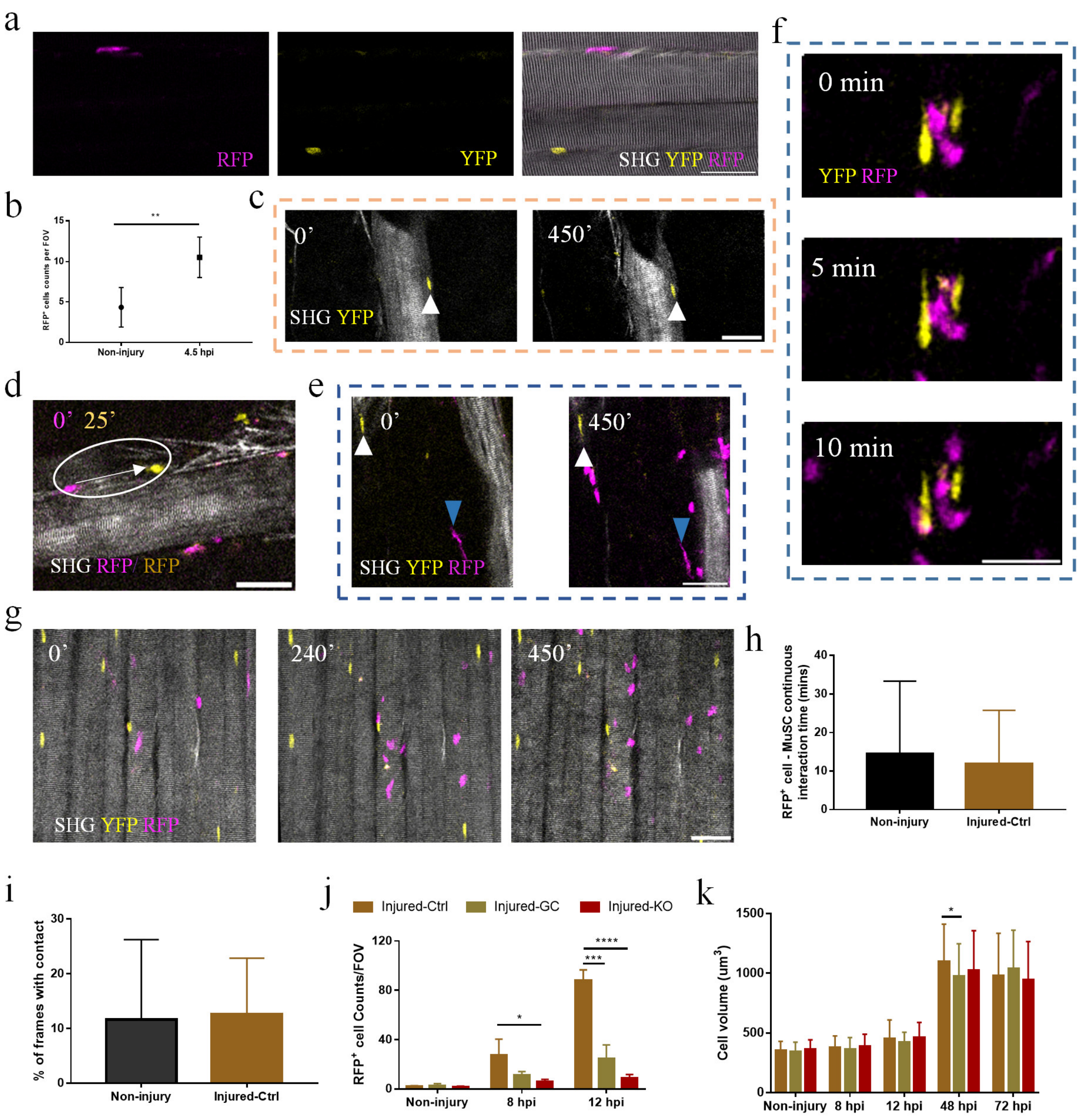


**Supplementary Fig. 4, related to Fig. 2: The dynamics of MuSCs and RFP$^+$ monocytes/macrophages immediately after injury.** (a) Representative images indicated distribution of the macrophages and MuSCs in the intact muscle fibers. Grey, SHG; yellow, YFP; magenta, RFP. (b) The RFP$^+$ cell density at non-injury and 4.5 hpi in the Injured-Ctrl group. (c) YFP$^+$ MuSCs at the beginning (left) and end (right) of 450 min time-lapse imaging starting at 4.5 hpi. Grey, SHG. Arrows: the traced MuSCs. (d) *In vivo* merged image of macrophages at 0 min (magenta) and 25 min (yellow) during time-lapse imaging. The circle and the arrow indicated the migration of the macrophage within 25 mins. Grey, SHG. (e) Image sequences indicated a motionless RFP$^+$ MO/MΦ (blue arrowheads) from the beginning (0 min) to the end (450 min), relative to YFP$^+$ MuSCs (white arrowheads) that did not change position. Grey, SHG; yellow, YFP; magenta, RFP. (f) Representative images of the interaction between RFP$^+$ MOs/MΦs and MuSCs within 10 minutes in the Injured-Ctrl group. (g) Maximum z intensity projections of TPEF image stacks of YFP$^+$ MuSCs (yellow), RFP$^+$ MOs/MΦs(magenta) at 0’, 240’ and 450’ post imaging in the non-injury group. The SHG (grey) was shown as an image for a certain depth rather than a projection. (h-i) The duration of the contact (h) and % of frames with contact (i) between MPs and RFP$^+$ MOs/MΦs from 4.5 to 12 hpi in the Non-injury and Injured-Ctrl groups. (j-k) The RFP$^+$ macrophage density (j) and MuSCs’ volume (k) at different time points of the Injured-Ctrl, Injured-GC, and Injured-KO groups. (n ≥ 3 mice per groups) Scale bars: 50 um in (a, c-e, g), 30 um in (f). Error bars: SD. (*: p<0.05; **: p<0.01; ***: p<0.001; ****: p<0.0001; student t-test). Source data are provided as a Source Data file.

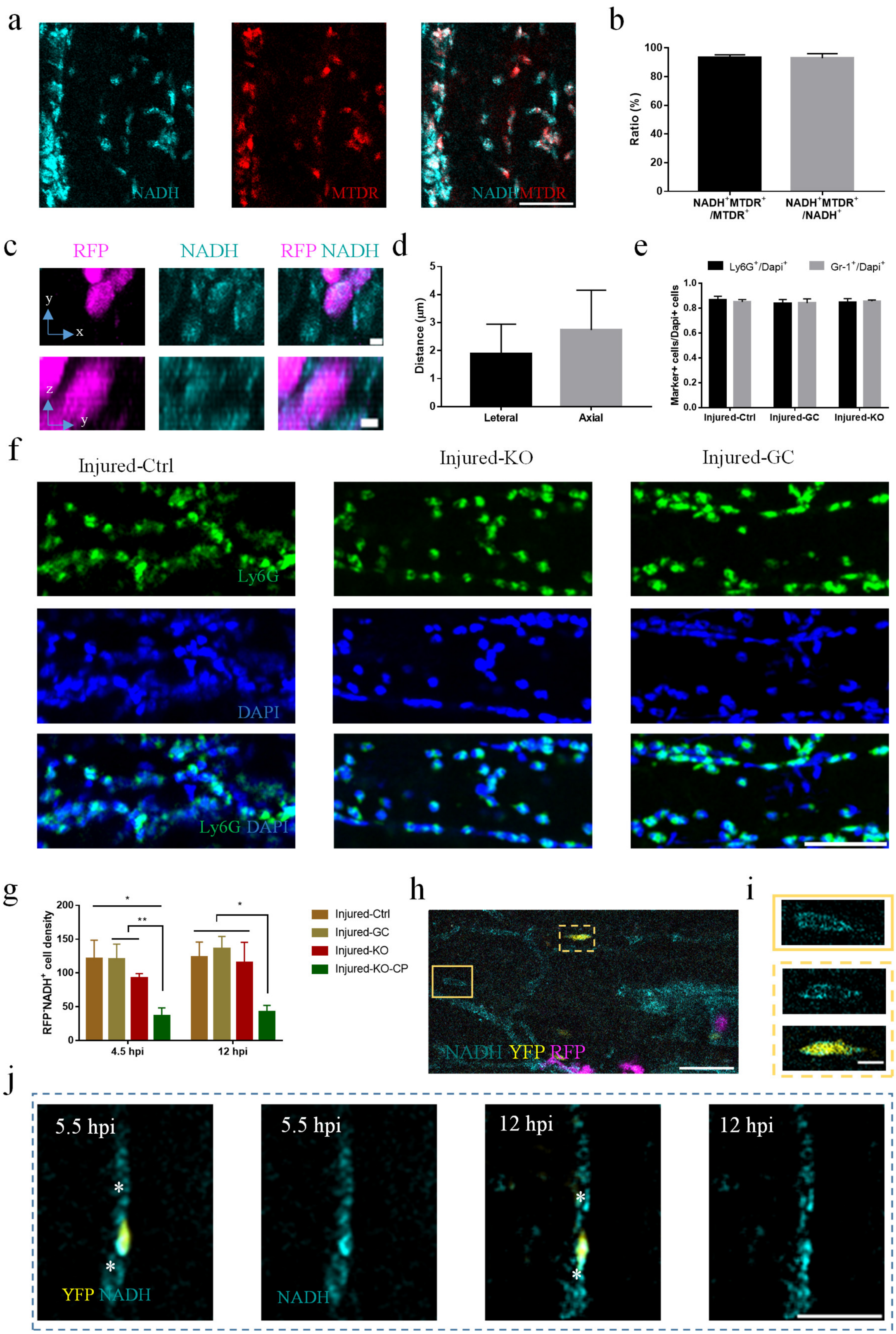
a
NADH
MTDR
NADH MTDR
b
Ratio (%)
100
80
60
40
20
0
NADH+MTDR+ /MTDR+
NADH+MTDR+ /NADH+
c
RFP
NADH
RFP NADH
y
x
z
y
d
Distance (μm)
5
4
3
2
1
0
Leteral
Axial
e
Ly6G+/Dapi+
Gr-1+/Dapi+
Marker+ cells/Dapi+ cells
1.0
0.8
0.6
0.4
0.2
0.0
Injured-Ctrl
Injured-GC
Injured-KO
f
Injured-Ctrl
Injured-KO
Injured-GC
Ly6G
DAPI
Ly6G DAPI
g
RFP-NADH+ cell density
200
150
100
50
0
*
**
*
4.5 hpi
12 hpi
Injured-Ctrl
Injured-GC
Injured-KO
Injured-KO-CP
h
NADH YFP RFP
i
j
5.5 hpi
5.5 hpi
12 hpi
12 hpi
YFP NADH
NADH

**Supplementary Fig. 5, related to Fig. 3: The interaction between MuSCs and $RFP^{-}NADH^{+}$ non-myogenic cells immediately after injury.** (a) Representative *in vivo* TPEF images of NADH (cyan), Mitotracker Deep Red (MTDR,green) for in-situ staining at 7 hpi. The merged image showed that most $NADH^{+}$ cells were stained by MTDR. (b) The statistics of overlap ratio of $NADH^{+}$ and $MTDR^{+}$ cells. (n=3 mice) (c) Representative images for the x-y and y-z projection of the macrophages in the RFP (Magenta) and NADH (Cyan) channels at 12 hpi. (d) The maximum distance of the cell boundary depicted by NADH signal to the RFP/YFP signal. (n =60 cells from 3 mice) (e)The ratio of Gr-$1^{+}$, $Ly6G^{+}$ cells at the injured site at 4.5 hpi in the Injured-Ctrl, Injured-KO, and Injured-GC mice. Most cells were Gr-$1^{+}$ or $Ly6G^{+}$ neutrophils. (f) Whole-mount TA muscle at 4.5 hpi in the Injured-Ctrl, Injured-KO, and Injured-GC mice stained with anti-ly6G (green) and Dapi (blue). (g) The $RFP^{-}NADH^{+}$ cell density at 4.5 and 12 hpi in the Injured-Ctrl, Injured-GC groups, Injured-KO, and Injured-KO-CP. ($n \geq 3$ mice per groups). (h) Maximum z intensity projections of TPEF image stacks of $YFP^{+}$ MuSCs (yellow), $RFP^{+}$ MOs/MΦs (magenta) and $NADH^{+}$ cells (cyan) at 4.5 hpi in the Injured-KO-CP. (i) The zoomed-in TPEF images of non-myogenic cells and $YFP^{+}$ MuSCs. Red, NADH; green, YFP. The imaging area corresponds to the orange solid and dash box region in (h). (j) Image sequences indicated that $YFP^{+}$ MuSCs established constant contact with two $RFP^{-}NADH^{+}$ non-myogenic cells in Injured-KO-CP during imaging. Yellow, YFP; magenta, RFP; cyan, NADH. Asterisks showed the $RFP^{-}NADH^{+}$ non-myogenic cells. Scale bars: 50 um in (a, f, h), 30 um in (j), 10 um in (i). 5 um in (c). Error bars: SD. (*: $p<0.05$; **: $p<0.01$; ***: $p<0.001$; ****: $p<0.0001$; student t-test). Source data are provided as a Source Data file.

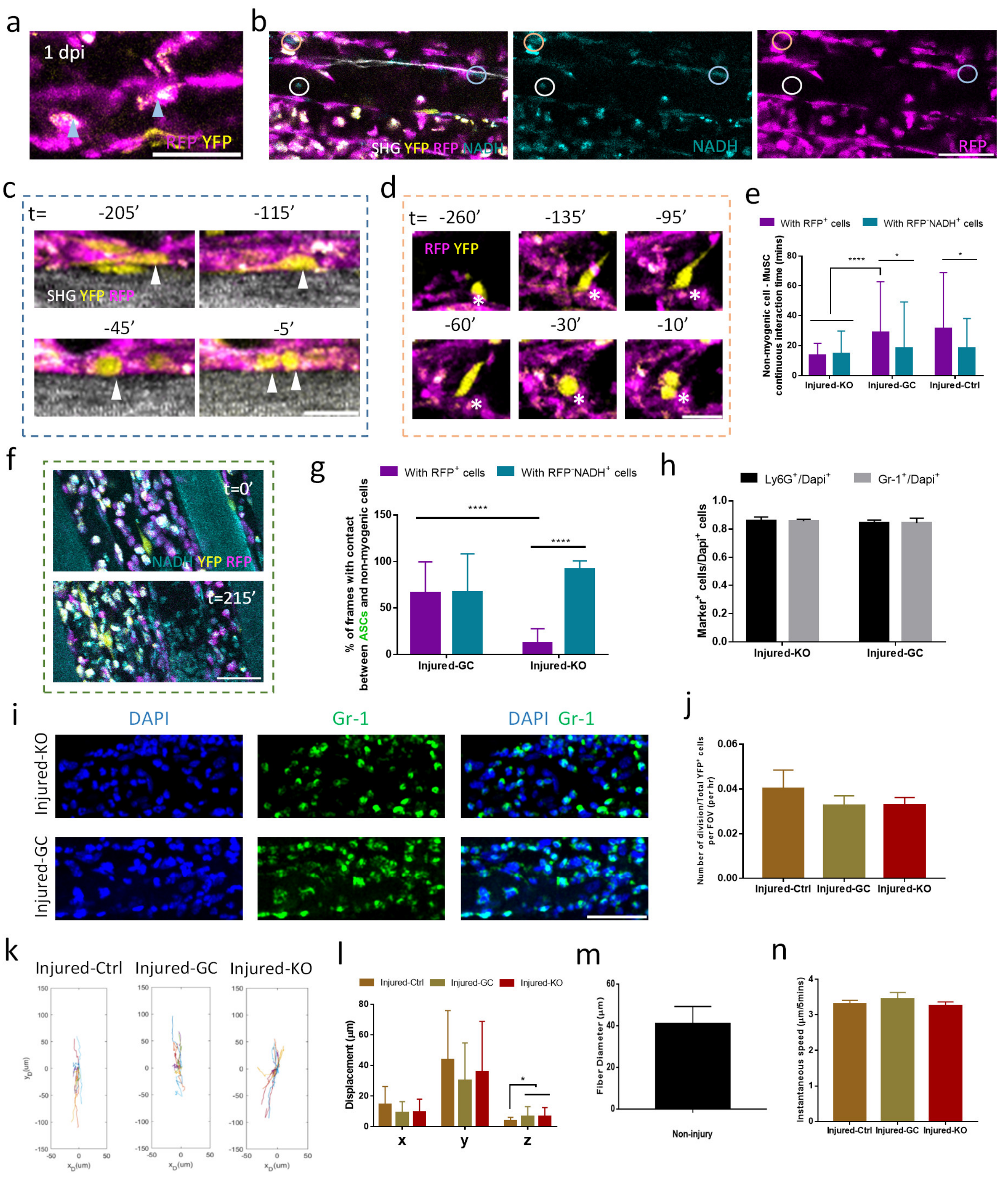

**Supplementary Fig. 6, related to Fig. 4 and 5: The spatiotemporal interaction between non-myogenic cells and activated satellite cells at the proliferation stage.** (a) Representative images of RFP$^+$ monocytes/macrophages (magenta) with strong autofluorescence (white) in the YFP channel (yellow), indicated by the arrowheads. (b) *In vivo* TPEF and SHG (grey) images of TA muscle at 1 dpi in Injured-Ctrl group. More than 90% NADH$^+$ (Cyan) cells were RFP$^+$ (magenta). RFP$^-$NADH$^+$ cells were indicated by the circles. (c) *In vivo* TPEF image sequences of YFP$^+$ ASCs (yellow) dividing parallel to the neighbor intact muscle fiber (SHG, grey) and surrounding by RFP$^+$ MOs/MΦs (magenta). Arrowheads indicated the traced ASCs. Daughter cells separated from each other at time 0. (d) Image sequence of the interaction between a traced ASC and RFP$^+$ MOs/MΦs at 2 dpi. Asterisks indicated a small region of RFP$^+$ MOs/MΦs that seemed contact ASC repetitively. (e) The duration of the contact between ASCs and non-myogenic cells at 1.5 dpi in the Injured-Ctrl, Injured-GC and Injure-KO groups. (f) Representative TPEF images indicated that the density of RFP$^-$NADH$^+$ non-myogenic cells increased during time-lapse image. Cyan, NADH; yellow, YFP; magenta, RFP. (g) The percentage of frames that ASCs contacted RFP$^+$, RFP$^-$NADH$^+$ cells during time-lapse imaging in the Injure-KO and Injured-GC groups. (h) The ratio of Gr-1$^+$, Ly6G$^+$ cells at the injured site at 1.5 dpi in Injure-KO and Injured-GC groups. Most cells were Ly6G$^+$/Gr-1$^+$ neutrophils. (n=3 mice). (i) Whole-mount TA muscle at 1.5 dpi in Injure-KO and Injured-GC groups stained with anti-Gr-1 (green) and Dapi (blue). Most cells were Gr-1$^+$. (j) The proliferation rate (Number of division/Total YFP$^+$ cells per FOV/hours) of ASCs at 1.5 dpi in the different groups (n≥3 mice). (k) Representative x-y trajectories (with t0 position as reference) of ASCs in the TA muscle at 1.5 dpi of three groups. Y-axis: the long axis of the muscle fiber. (l) The displacement in x, y, z-axis of ASCs within two hours at 1.5 dpi in three groups. (n ≥ 24 cells from more than three mice per group). (m) Muscle fiber diameter measured in non-injured TA muscle based on SHG signal. 39 muscle fibers from 3 mice. (n)The instantaneous speed of ASCs within two hours at 1.5 dpi in three groups. (n ≥ 24 cells from more than three mice per group). Scale bars, 50 um in (a-b), (f), (i); 30 um in (c-d). Error bars: SD. (*: p<0.05; **: p<0.01; ***: p<0.001; ****: p<0.0001; student t-test). Source data are provided as a Source Data file.

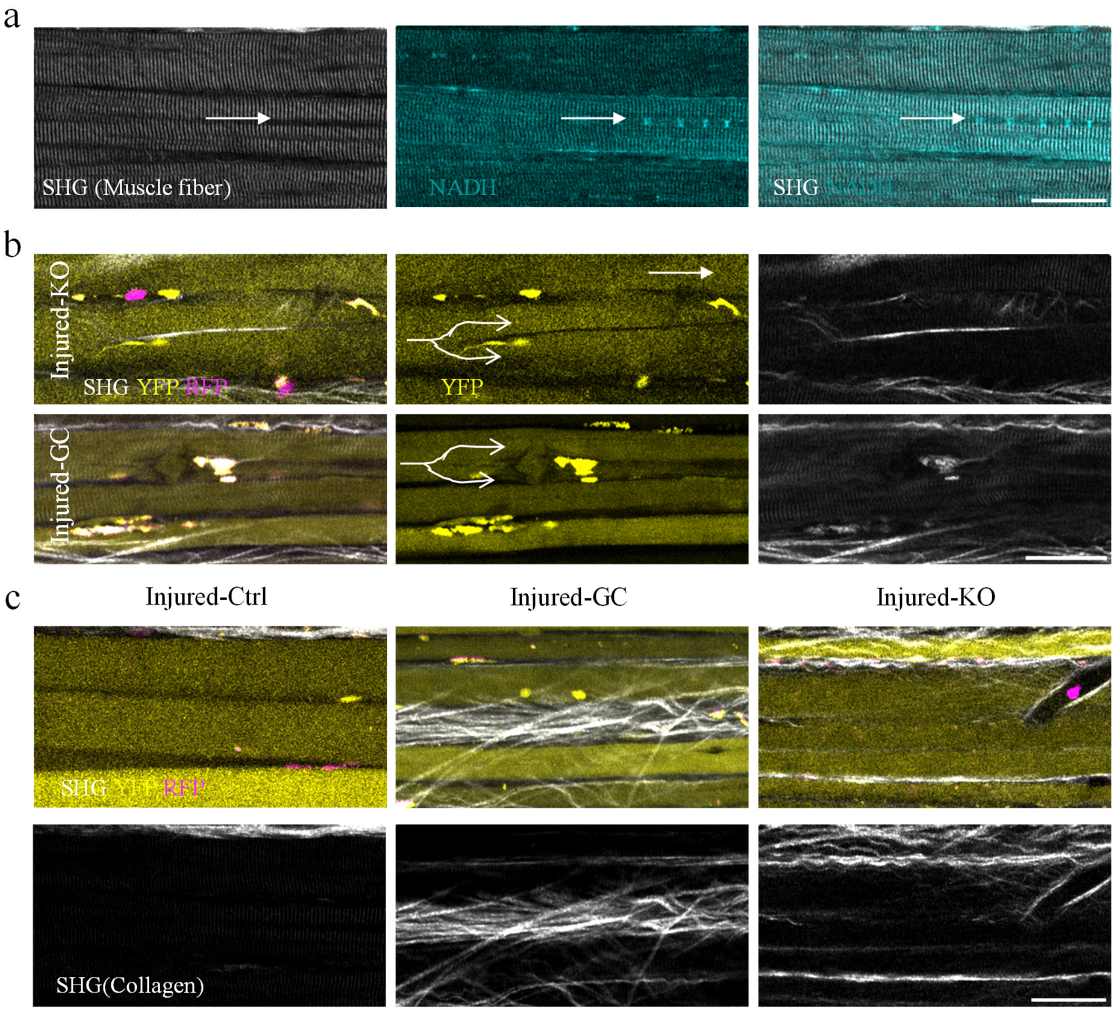


**Supplementary Fig. 7, related to Fig. 6: The depletion of macrophages impaired the muscle regeneration.** (a) Represent SHG (grey) and NADH (cyan) image of regenerated muscle fiber at 15 dpi in Injured-Ctrl groups. Arrow: a dark line at the center of the SHG signal of the muscle fiber and centrally-aligned nuclei in the muscle fiber indicated by NADH. (b) Represent TPEF and SHG (grey) images of regenerated muscle fiber (YFP, yellow) at 15 dpi in the Injured-KO and Injured-GC groups. The lines and arrows indicated the bifurcated muscle fibers. (c) Representative TPEF and SHG images to indicate the msucular microenvironment at 15 dpi in the Injured-Ctrl, Injured-GC, and Injured-KO. Top: merged images, magenta, RFP; yellow, YFP; grey, SHG. Bottom: enhanced SHG signal of collagen. Scale bars: 50 um.

### Supplementary Videos

**Supplementary Video 1: The time-lapse imaging indicated the infiltration of $RFP^+$ monocytes/macrophages and $RFP^-$ non-myogenic cells from 4.5 to 12 hpi in the Injured-Ctrl groups.** $YFP^+$ MuSCs were motionless but non-myogenic cells were highly dynamic. Movie frames are Z stack maximal projections comprised of 20 slices spaced 3 um and captured at 5 mins interval for 450 mins. The image started at 4.5 hpi. Cyan, NADH; yellow, YFP; magenta, RFP. Scale bar: 50 um.

**Supplementary Video 2: Dynamics of a dwelling $RFP^+$ monocytes/macrophages from 4.5 to 12 hpi in the Injured-Ctrl groups.** One of the $RFP^+$ cells dwelled in the injured site during imaging. Movie frames are Z stack maximal projections comprised of 6-8 slices spaced 3 um and captured at 5 mins interval for 450 mins. The image started at 4.5 hpi. Magenta, RFP; grey, SHG. Scale bar: 50 um.

**Supplementary Video 3: Direct interaction between MuSCs and non-myogenic cells during time-lapse imaging in the Injured-Ctrl groups.** MuSCs established transient contact with different non-myogenic cells frequently in the Injured-Ctrl groups. Movie frames are Z stack maximal projections comprised of 6-8 slices spaced 3 um and captured at 5mins intervals for 450 mins. The image started at 4.5 hpi. Cyan, NADH; yellow, YFP; magenta, RFP. Scale bar: 30 um.

**Supplementary Video 4: The time-lapse imaging indicated the infiltration of $RFP^+$ monocytes/macrophages from 4.5 to 12 hpi in the Non-injury groups**. $YFP^+$ MuSCs were motionless but $RFP^+$ cells were highly dynamic. Movie frames are Z stack maximal projections comprised of 20 slices spaced 3 um and captured at 5 mins interval for 450 mins. The image started at 4.5 hpi. Yellow, YFP; magenta, RFP; grey, SHG. Scale bar: 50 um.

**Supplementary Video 5: The time-lapse imaging indicated the infiltration of $RFP^+$ monocytes/macrophages and $RFP^-$ non-myogenic cells from 4.5 to 12 hpi in the Injured-GC groups.** $YFP^+$ MuSCs were motionless but non-myogenic cells were highly dynamic. Only few $RFP^+$ cells were recruited to the injured site. Movie frames are Z stack maximal projections comprised of 20 slices spaced 3 um and captured at 5 mins interval for 450 mins. The image started at 4.5 hpi. Cyan, NADH; yellow, YFP; magenta, RFP. Scale bar: 50um**.**

**Supplementary Video 6: The time-lapse imaging indicated the infiltration of $RFP^+$ monocytes/macrophages and $RFP^-$ non-myogenic cells from 4.5 to 12 hpi in the Injured-KO groups.** $YFP^+$ MuSCs were motionless but non-myogenic cells were highly dynamic. Only few $RFP^+$ cells were recruited to the injured site. Movie frames are Z stack maximal projections comprised of 20 slices spaced 3 um and captured at 5 mins interval for 450 mins. The image started at 4.5 hpi. Cyan, NADH; yellow, YFP; magenta, RFP. Scale bar: 50um**.**

**Supplementary Video 7: Direct interaction between MuSCs and $RFP^-NADH^+$ non-myogenic cells during time-lapse imaging in the Injured-GC groups.** MuSCs established transient contact with different non-myogenic cells frequently in the Injured-GC groups. Movie frames are Z stack maximal projections comprised of 6-8 slices spaced 3 um and captured at 5mins intervals. The image started at 4.5 hpi. Cyan, NADH; yellow, YFP; magenta, RFP. Scale bar: 30um.

**Supplementary Video 8: Direct interaction between MuSCs and $RFP^-NADH^+$ non-myogenic cells during time-lapse imaging in the Injured-KO groups.** MuSCs established transient contact with different non-myogenic cells frequently in the Injured-KO groups. Movie frames are Z stack maximal projections comprised of 6-8 slices spaced 3 um and captured at 5mins intervals. The image started at 4.5 hpi. Cyan, NADH; yellow, YFP; magenta, RFP. Scale bar: 30um.

**Supplementary Video 9: The time-lapse imaging indicated the infiltration of $RFP^+$ monocytes/macrophages and $RFP^-$ non-myogenic cells from 4.5 to 12 hpi in the Injured-KO-CP groups.** $YFP^+$ MuSCs were motionless but non-myogenic cells were highly dynamic. Movie frames are Z stack maximal projections comprised of 20 slices spaced 3 um and captured at 5 mins interval for 450 mins. The image started at 4.5 hpi. Cyan, NADH; yellow, YFP; magenta, RFP. Scale bar: 50um**.**

**Supplementary Video 10: Direct interaction between MuSCs and $RFP^-NADH^+$ non-myogenic cells during time-lapse imaging in the Injured-KO-CP groups.** MuSCs established transient contact with different non-myogenic cells frequently in the Injured-KO-CP groups. Movie frames are Z stack maximal projections comprised of 6-8 slices spaced 3 um and captured at 5 mins intervals. The image started at 4.5 hpi. Cyan, NADH; yellow, YFP; magenta, RFP. Scale bar: 30 um.

**Supplementary Video 11a: $YFP^+$ ASCs migrated along the longitudinal axis of the neighbor intact muscle fiber and interacted with $RFP^+$ macrophages before division at 2 dpi in the Injured-Ctrl groups.** Time-lapse *in vivo* imaging of $YFP^+$ ASC (yellow) migrating parallel to the neighbor intact muscle fiber (grey) and interacting with different $RFP^+$ macrophages (magenta) at 2 dpi in the Injured-Ctrl group. Movie frames are Z stack maximal projections comprised of 6-8 slices spaced 3 um and captured at 5 mins interval. Blue points indicate the traced $YFP^+$ ASC. Scale bar: 50 um.

**Supplementary Video 11b: $YFP^+$ ASCs interacted with $RFP^+$ monocytes/macrophages before division at 2 dpi in the Injured-Ctrl groups.** Movie frames are Z stack maximal projections comprised of 6-8 slices spaced 3 um and captured at 5mins interval. Yellow, YFP; magenta, RFP. Scale bar: 50 um.

**Supplementary Video 12a: $YFP^+$ ASCs migrated along the longitudinal axis of the neighbor intact muscle fiber and interacted with non-myogenic cells at 1 dpi in the Injured-Ctrl groups.** Time-*lapse in vivo* imaging of $YFP^+$ ASC (yellow) migrating parallel to the neighbor muscle fiber and interacting with different $RFP^+$ macrophages (magenta), $RFP^-NADH^+$ cells (cyan) at 1 dpi in the Injured-Ctrl groups. Movie frames are Z stack maximal projections comprised of 6-8 slices spaced 3 um and captured at 5 mins intervals. Cyan, NADH; yellow, YFP; magenta, RFP. Scale bar: 50 um.

**Supplementary Video 12b: $YFP^+$ ASCs interacted with $RFP^+/RFP^-$ non-myogenic cells before division at 1.5 dpi in the Injured-Ctrl groups.** Movie frames are Z stack maximal projections comprised of 6-8 slices spaced 3 um and captured at 5 mins interval. Blue points indicate the traced $YFP^+$ ACSs. Cyan, NADH; yellow, YFP; magenta, RFP. Scale bar: 50 um.

**Supplementary Video 12c: $YFP^+$ ASCs migrated along the longitudinal axis of the neighbor intact muscle fiber and interacted with non-myogenic cells before division at 1.5 dpi in the Injured-Ctrl groups.** Time-lapse *in vivo* imaging of $YFP^+$ ASC (yellow) interacting with different $RFP^+$ macrophages (magenta), $RFP^-NADH^+$ cells (cyan) at 1.5 dpi in the Injured-Ctrl group. Movie frames are Z stack maximal projections comprised of 6-8 slices spaced 3 um and captured at 5 mins intervals. Scale bar: 50 um.

**Supplementary Video 13: $YFP^+$ ASCs migrated along the longitudinal axis of the neighbor muscle fiber and interacted with $RFP^+/RFP^-$ $NADH^+$ non-myogenic cells before division at 1.5 dpi in the Injured-KO groups.** Time-lapse *in vivo* imaging of $YFP^+$ ASCs (yellow) migrating parallel to the neighbor muscle fiber and interacting with different $RFP^+$ macrophages (magenta) and $RFP^-$ $NADH^+$ non-myogenic cells (cyan) at 1.5 dpi in the Injured-KO groups. Movie frames are Z stack maximal projections comprised of 6-8 slices spaced 3 um and captured at 5 mins intervals for 160 mins. Scale bar: 50 um.

**Supplementary Video 14a and S14b: $YFP^+$ ASCs migrated along the longitudinal axis of the neighbor muscle fiber and interacted with $RFP^+/RFP^-$ $NADH^+$ non-myogenic cells before division at 1.5 dpi in the Injured-GC groups.** Time-lapse *in vivo* imaging of $YFP^+$ ASCs (yellow) migrating parallel to the neighbor muscle fiber and interacting with different $RFP^+$ macrophages (magenta) and $RFP^-NADH^+$ non-myogenic cells (cyan) at 1.5 dpi in the Injured-GC group. Movie frames are Z stack maximal projections comprised of 6-8 slices spaced 3 um and captured at 5 mins interval. Scale bar: 50 um.
